# Structure-Based Discovery of GTP-Enhanced Rab10 Modulators and Their Effects on Alzheimer’s Disease-Relevant Cellular Phenotypes

**DOI:** 10.64898/2026.10.30.763175

**Authors:** Satyaveni Malasala, Nathan Cox, Grady Sellman, Yan-Hua Chen, Qun Lu

## Abstract

Rab10-mediated membrane trafficking has been implicated in Alzheimer’s disease (AD), but pharmacological tools for investigating its function remain limited. Here, we screened more than 80 million compounds using structure-based virtual screening and prioritized five candidate Rab10 modulators (**R1-R5**). These compounds exhibited favorable docking scores, stable modeled protein-ligand complexes in molecular dynamics (MD) simulations and predicted acceptable ADME/toxicity profiles. Comparative docking against Rab5, Rab8, and Rab11 predicted preferential interactions with GTP-associated Rab10. **R1** and **R2** yielded the most favorable Rab10 scores, whereas **R3** showed the largest differences between GTP-associated, GDP-containing, and nucleotide-free Rab10 models. Surface plasmon resonance (SPR) revealed micromolar apparent affinities of **R1-R4** without added GTP and nanomolar apparent affinities under GTP-containing conditions, generally favoring GTP-bound Rab10 over Rab5. In a cell-based GTP quantification assay, **R1**, **R2**, **R3**, and **R5** reduced GTP-bound Rab10, whereas **R4** increased it. **R3** produced dose- and time-dependent reductions in GTP-bound Rab10 and was selected for functional studies. **R3** altered endosomal organization and reduced EEA1 and LAMP1 expressions. In amyloid precursor protein (APP)/presenilin-1-expressing CHO cells, **R3** redistributed intracellular APP, reduced APP carboxyl-terminal fragments (APP-CTFs), and decreased enlarged endosome-like structures. In neuronal differentiated PC12 cells, **R1**-**R3** attenuated NGF withdrawal-induced cytoskeletal disruption and cell death, accompanied by reduced APP-CTF accumulation and tau phosphorylation. These studies highlight novel Rab10 modulators and support their further validation as candidate pharmacological probes for investigating Rab10-associated trafficking and AD-relevant cellular phenotypes.

## Introduction

Alzheimer’s disease (AD) is a complex neurodegenerative disorder in which amyloid-β (Aβ) accumulation, tau pathology, neuroinflammation, oxidative stress, and disruption of intracellular membrane trafficking contribute to disease initiation and progression. Importantly, a substantial proportion of older individuals remain cognitively resilient despite meeting neuropathological criteria for AD at autopsy, including the presence of senile plaques and neurofibrillary tangles^1–6^. This clinicopathological dissociation highlights the need to identify early, modifiable cellular processes that influence disease vulnerability and resilience. Accordingly, therapeutic strategies are expanding beyond the removal of Aβ and tau aggregates to include correction of early pathogenic mechanisms, particularly endolysosomal trafficking, autophagy, and cellular stress responses^7^.

Rab proteins are small guanosine triphosphatases that act as central regulators of vesicle formation, cargo sorting, membrane identity, and intracellular trafficking^8–10^. Among these proteins, Rab10 has emerged as a compelling AD-relevant target because of its established roles in endosomal trafficking, recycling, autophagy, and cargo delivery to lysosomal compartments^11–14^. These pathways are strongly implicated in AD, where early endosomal abnormalities, defective autophagic clearance, and altered amyloid precursor protein (APP) trafficking are prominent pathological features. Dysregulated Rab10 activity may therefore directly influence the trafficking and processing of APP, endolysosomal homeostasis, and neuronal survival.

Multiple lines of evidence support an important involvement of Rab10 in AD. Rab10 has been identified among resilience-associated genes in human genomic studies^15^, and a rare RAB10 variant (rs142787485) has been associated with reduced AD risk and cognitive resilience in individuals with high genetic predisposition to disease^16–18^. Elevated Rab10 expression and phosphorylation have also been reported in AD brains compared with healthy controls^19–21^. Functionally, Rab10 knockdown reduces Aβ42 production and lowers the Aβ42/Aβ40 ratio, whereas Rab10 overexpression enhances amyloidogenic APP processing^15,22,23^. Together, these findings position Rab10 as an active regulator of AD-related APP metabolism and a potential determinant of disease susceptibility.

Rab10 is also relevant to other neurodegenerative disorders. In Parkinson’s disease (PD), Rab10 phosphorylation at threonine 73 is regulated by leucine-rich repeat kinase 2 (LRRK2), mutations of which are the most common genetic cause of late-onset familial and sporadic PD, linking Rab10 to LRRK2-dependent vesicular trafficking and lysosomal dysfunction^24–27^. Because impaired trafficking and autophagy are shared features across neurodegenerative diseases, direct pharmacological modulation of Rab10 may provide a strategy to restore cellular homeostasis across multiple pathological settings.

Despite the biological rationale, direct and selective pharmacological tools for Rab10 remain scarce. Reported approaches include the natural product amentoflavone, which has been proposed as a Rab10 inhibitor based primarily on computational binding analysis, and indirect inhibition of Rab10 phosphorylation through LRRK2 inhibitors such as PF-06447475^28–30^. Genetic suppression of RAB10 is also feasible, but it does not provide the temporal control, reversibility, or translational potential of small-molecule modulation. Thus, identifying direct, cell-permeable Rab10 small-molecule modulators (SMMs) remains an important unmet need.

Here, we investigated Rab10 as a candidate pharmacological target in AD-related trafficking dysfunction. Structure-based virtual screening of more than 80 million compounds identified five candidate SMMs, **R1-R5**, with favorable docking scores and stable modeled protein-ligand complexes in molecular dynamics (MD) simulations. Comparative docking against Rab5, Rab8, and Rab11 predicted preferential interactions with GTP-associated Rab10. Surface plasmon resonance (SPR) analysis demonstrated GTP-associated enhancement of apparent Rab10 binding. Cell-based GTP quantification identified inhibitor-like effects for **R1, R2, R3, and R5** and an activator-like effect for **R4**. Compound treatment altered endolysosomal organization, improved neuronal survival in differentiated PC12 cells, and modified APP-related cellular phenotypes.

## Materials and Methods

### Reagents and Cell Lines

Recombinant Rab10 protein was obtained from Creative Biomart (NY, USA). Small molecule compounds **R1, R2** and **R3** were from Chembridge whereas **R4** was obtained from Enamine. **R5** was supplied by the National Cancer Institute Experimental Therapeutics Program. Lipopolysaccharide and insulin were from Sigma Aldrich and Invitrogen, respectively. HEK293 cells were obtained from American Type Culture Collection (ATCC) (Manassas, VA) and cultured as described. Chinese Hamster Ovary (CHO) cells (PS70) overexpressing presenilin 1 and APP were a gift from Dr. Weiming Xia. Compounds were solubilized in 5% DMSO (Sigma Aldrich).

### In Silico Screening Ligand Preparation

The chemical libraries were derived from Mcule and ZINC for high throughput screening. The LigPrep module of Schrödinger software was used to prepare, neutralize, desalt, and adjust tautomeric states of the library. Then, OPLS4 force field was employed to minimize their energy (LigPrep, Schrödinger, NY).

### Molecular Docking

All computational studies were carried out by the Schrödinger version 2024-02. Structure- and ligand-based modelling as well as MD simulations were performed using Desmond version 2024-02 of Schrödinger Small Molecule Suite. Briefly, the crystallographic structure of Rab10 (PDB code: 5SZJ) was downloaded from Protein Data Bank at RCSB (https://www.rcsb.org/) and prepared using the protein preparation wizard module of the Schrödinger software package. After the introduction of hydrogen bonds, missing side chains and loops were filled. Subsequently, the OPLS4 force field was employed for optimization and minimization. The receptor grid generation module was generated around the GTP binding pocket. The docking studies were conducted using the Glide module of Schrödinger software. Initially, the Glide Extra Precision (G-XP) mode was used for predicting the binding mode of HM compounds, and the top-ranked output was analyzed.

### MD Simulation

MD simulations were conducted to evaluate the effectiveness of the compounds screened through molecular docking. The simulations were performed using the Desmond module of the Schrödinger Suite (v2024-02). The system was set up with the TIP3P water model as the solvent, and the simulations were carried out under NPT ensemble conditions with a simulation time of 100 ns, a temperature of 300 K, and a relaxation time of 1 ns. The OPLS_2004 force field was applied throughout the simulations. Water molecules were explicitly represented using the TIP3P model. The TIP3P water model was configured in an orthorhombic box with periodic boundary conditions of 15 Å × 15 Å × 15 Å along the x, y, and z axes, centered on the protein-ligand complex. The Coulomb interactions were calculated with a cutoff radius of 9.0 Å. Pressure and temperature were controlled using the Martyna-Tuckerman-Klein chain coupling scheme (with a coupling constant of 2.0 ps) and the Nose-Hoover chain coupling scheme, respectively. Nonbonded forces were computed using an r-RESPA integrator, with short-range forces updated every step and long-range forces updated every three steps. Trajectories were recorded at intervals of 100 ps for subsequent analysis. Protein-ligand interactions were analyzed using the Simulation Interaction Diagram tool within the Desmond MD package. The stability of the simulations was assessed by monitoring the root-mean-square deviation (RMSD) of the ligand and protein atom positions over time.

### Antibodies

Primary antibodies listed were as follows: rabbit anti-EEA1 Antibody (Dilution 1:70 for immunostaining and 1:1000 for Western blot, sigma; Cat no: 07-1820); pRab10 (Dilution 1:500 for Western blot; 1:250 for immunofluorescence (IF) staining; Abcam; Cat no: ab241060); Rab10 (Dilution 1:500 for Western blot; 1:100 dilution for immunohistochemical (IHC) staining; 1:250 for IF; 1:1000 for co-IP; Proteintech; Cat no: 11808-1-AP); LAMP1 (Dilution 1:500 for Western blot; Proteintech; Cat no: 21997-1-AP); GM130 (Dilution 1:1000 for Western blot; BD transduction laboratories; Cat no: 560257); Histone H3 (Dilution 1:1000 for Western blot; Cell signaling technology; Cat no: #9715); 22C11 anti-APP (Dilution 1:1000 for Western blot; 1:500 for immunoprecipitation (IP); ThermoFisher, Catalog # 13-9749-82) and CT695 anti-APP CTF (Dilution 1:2000 for Western blot; 1:1000-2500 for IP; ThermoFisher, Cat # 51-2700); mouse anti-tau (PHF-1) (Dilution 1:1000 for Western blot; gift from Dr. Peter Davies). For immunofluorescent labeling, Alexa Fluor™ 488 (Dilution 1:600, Invitrogen; Cat: A-11008) and Cy3 were used (Dilution 1:600; Jackson Immuno Research Inc.; Cat: 111-165-003). For Western blot, mouse anti-HRP conjugated antibody (Dilution 1:2500, Invitrogen; Cat: # A-10651) and rabbit anti-HRP conjugated antibody were used (Dilution 1:2500, Invitrogen; Cat: # 31460).

### Immunofluorescent Light Microscopy

HEK293 cell cultures were maintained until they reached the required confluency for splitting. Cells were treated with the corresponding small molecule compounds for 2 h. After treatment, cells on coverslips were fixed in methanol at -20°C. After rinsing the cells with phosphate-buffered saline (PBS), the cells were blocked in 5% BSA in PBS. The primary antibody incubation was followed by secondary antibody treatment. Finally, the coverslips with cells were mounted using Fluoro-Gel II with DAPI (Cat# 1785-50 EMS, PA) for microscopic analysis. They were imaged using a Zeiss Axio Observer 7 Imager Fluorescence Microscope equipped with an AxioCam camera (Carl Zeiss, Thornwood, NY).

### Image Quantification

ImageJ was used to measure the fluorescence signal, as described, as the colored image was converted into a 16-bit grayscale image, and an additional copy was created. The region of interest (ROI) was selected, and the threshold for the ROI was adjusted. The subtract background function was applied to minimize background autofluorescence. After selecting the ROI, a binary image in black and white was generated. This binary image was then overlaid onto the second grayscale image as a reference. Finally, the binary image containing the ROI was used to quantify the fluorescence intensity, expressed as mean pixel intensity.

### SDS-PAGE and Western Blotting

Proteins were resolved on 8-16% and 10-20% gradient Tris-glycine gels (Thermo Fisher Scientific, cat # XP10200BOX). The proteins on the gels were transferred to nitrocellulose membranes (Biorad, cat #170415) by semi-dry trans-Blot Turbo transfer system (Biorad). The membranes were blocked with 5% milk and then incubated with primary antibodies. Afterwards, the membranes were washed with 5% milk followed by incubation with secondary antibodies, as previously stated. The blots were washed in 0.1% TBST and scanned on a Clarity Western ECL Substrate. Quantitation of bands was performed using ChemiDOC (Bio-Rad). Blots were probed for GAPDH and Histone H3 as housekeeping proteins to ensure equal loading of samples.

### SPR Validation

SPR experiments were performed using a Biacore 1S+ instrument equipped with a CM5 sensor chip. All solutions were prepared with Milli-Q water and filtered through a 0.22-μm membrane. The sensor surface was activated with EDC/NHS at a flow rate of 10 μL/min for 200 s. Rab10 (100 μg/mL in 10 mM sodium acetate, pH 5.0) was then injected at 5 μL/min for 500 s and covalently immobilized to a level of approximately 5,000 response units (RU). Unbound protein was removed by washing with running buffer. Compounds **R1–R5** were prepared at concentrations ranging from 0.0136 to 10 μM in 20 mM Tris buffer (pH 7.4) containing 5% DMSO and injected at 10 μL/min for 150 s in the presence or absence of GTP. Sensorgrams were analyzed using Biacore evaluation software and GraphPad Prism.

### Rab10-CGQ-ELISA Activation Assay

The Rab10 Cell-based GTP Quantification Enzyme-linked immunosorbent assay (Rab10-CGQ-ELISA activation assay) measures the level of GTP-bound Rab10 protein in cell lysates. HEK293 cells (70% confluent) were treated with different concentrations (0.05, 0.5, 5, 50 and 100 µM) of R1-R5 compounds. For time-dependent Rab10-CGQ-ELISA, **R1-R5** were tested for time intervals including 0 min, 15 min, 30 min, 1 hour (h), 2 h, and 4 h. The cells were lysed and ∼ 0.25 mg/mL protein was used for Rab10-CGQ-ELISA analysis. After placing the collected lysates in a 96-well Rab10-GTP binding plate, free or protein-bound GTP was washed out and the Rab10 bound GTP levels were measured on Biotek Synergy HT (Agilent, Winooski, VT).

### PC12 Cell Culture and Differentiation

PC12 cells were seeded onto poly-L-lysine-coated 6-well plates. Cells were initially maintained in DMEM 11960 growth medium supplemented with 1% L-glutamine, 1% penicillin-streptomycin, 5% fetal bovine serum (FBS), and 10% horse serum. After the initial attachment period, five of the six wells were treated with nerve growth factor (NGF, 50 ng/mL) and maintained under this condition for 10 days, with medium changes performed every 2–3 days. These NGF-treated cells were designated as neuronal differentiated PC12 cells.

After 10 days of NGF exposure, four of the five NGF-treated wells were subjected to NGF withdrawal by replacing the medium with serum-free PC12 DMEM 11960 medium containing 1% L-glutamine and 1% penicillin-streptomycin. The experimental conditions were assigned as follows: NGF+ (neuronal differentiated PC12 cells maintained in medium with 50 ng/mL NGF), NGF− (neuronal differentiated PC12 cells maintained in medium without NGF), **R1** 50 μM (NGF withdrawal plus **R1** at 50 μM), **R2** 50 μM (NGF withdrawal plus **R2** at 50 μM), and **R3** 50 μM (NGF withdrawal plus **R3** at 50 μM). Following the treatment, cells were processed for immunofluorescence staining and/or survival analysis.

### Statistical analysis

Statistical analyses were performed using GraphPad Prism version 10.0.2(232) (La Jolla, CA). For immunostaining studies, data are expressed as mean ± SEM from at least three independent experiments. A p-value of <0.05 was considered statistically significant. Differences among the top five small molecules were assessed using one-way ANOVA.

## Results

### *In silico* Identification of Small Molecule Modulators with Rab10 Binding

Using structure-based rational design, we sought to identify small molecule modulators (SMMs) of Rab10 protein (**Figure 1A**). We performed computer-aided high-throughput, *in silico* screening over 80 million structurally diverse chemical compounds from various databases, including Mcule and ZINC (**Figure 1**). We selected the top 100 hits based on optimal docking scores and strong protein-ligand interactions (**Supplementary Table S1)**.

**Figure 1.**
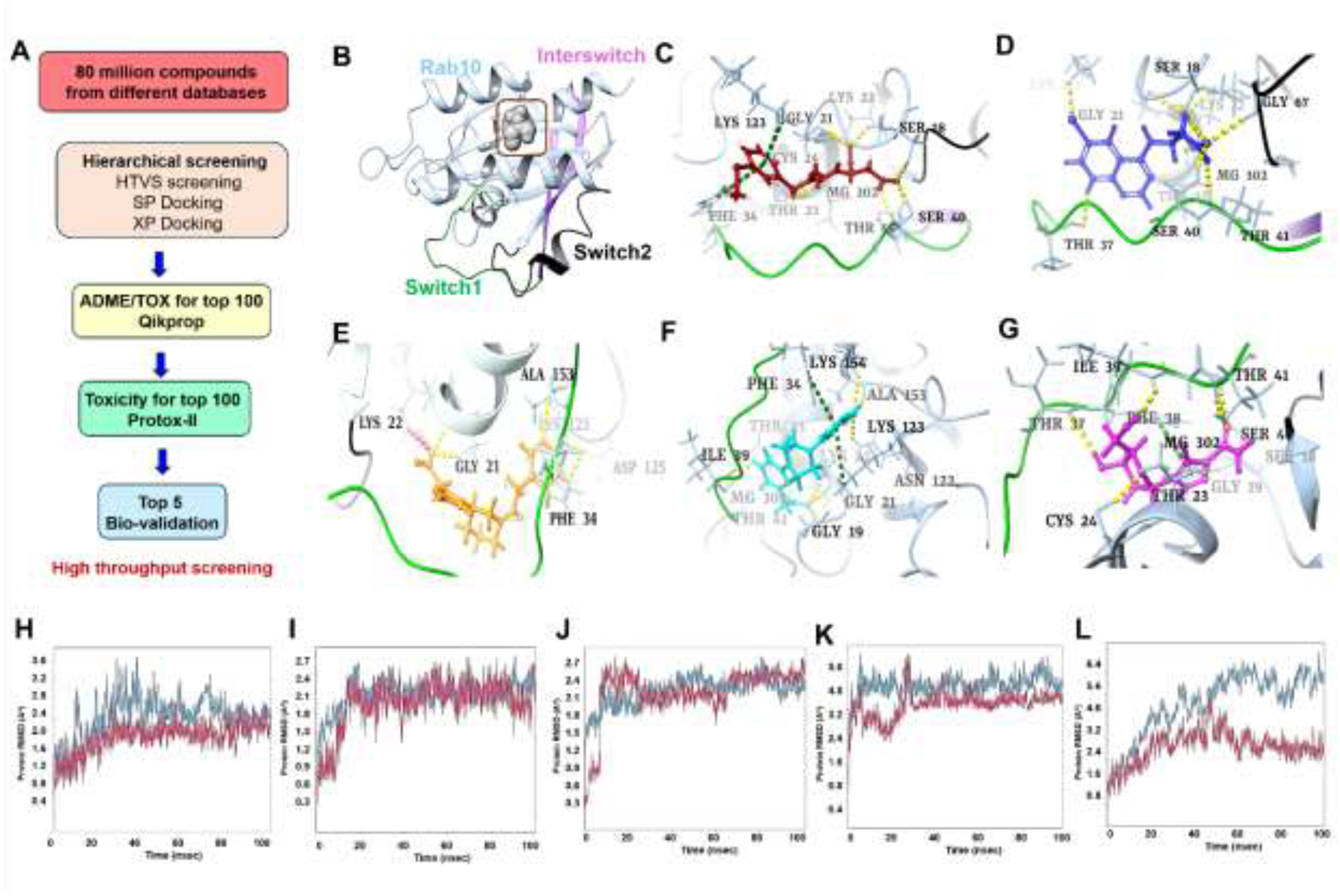
Structural and molecular dynamic analysis of Rab10 and its interactions with Rab10 SMMs R1-R5. **(A)** Schematic workflow of CADD and *in silico* high throughput screening to identify Rab10 modulators. **(B)** Overview of the Rab10 protein (light blue), with the binding site illustrated in 3D (gray, CPK representation). The Switch I region (amino acids 31-43) is highlighted in green, the Switch II region (amino acids 66-82) in black, and the inter switch region (amino acids 44-63) in magenta (PDB ID: 5SZJ). **(C-G)** 3D views of the Rab10 GTP-binding site showing interactions of active site residues with compounds **R1 (C), R2 (D), R3 (E), R4 (F), and R5 (G)**. Hydrogen bonds are represented by yellow dashed lines, and π–π interactions are shown in green. Binding site residues are color-coded based on interaction type: hydrophobic (green), polar (cyan), positively charged (violet), and negatively charged (red). **(H-L)** Protein-ligand RMSD plots for Rab10 complexes with compounds **R1 (H), R2 (I), R3 (J), R4 (K), and R5 (L)** were generated from 100 ns MD simulations using the Desmond module of Schrödinger software. The Y-axis represents RMSD values in angstroms (Å), while the X-axis denotes simulation time in nanoseconds (ns). The calculated RMSD values for all five compounds remained below 5 Å.

The MM/GBSA binding energy and molecular docking analysis showed that the top five ranking SMMs **R1-R5** displayed different preferences in Rab10 binding regarding the presence or absence of GTP within the binding site (holo form-**Supplementary Table S2** and apo form**-Supplementary Table S3**). Among the selected SMMs, **R3** exhibited an optimal binding with a binding energy of -43.21 Kcal/mol in the GTP-bound form and -45.14 Kcal/mol in the apo form, which is similar or even higher than GTP. The other SMMs (e.g., **R1, R2,** and **R5**) also demonstrated binding, albeit weaker, particularly in the absence of GTP. The SMM **R4**, though showing tight binding when GTP-bound, displayed a dramatic reduction of binding affinity in the apo form. These data indicated that the potential optimal binding between **R4** and Rab10 is dependent on GTP-induced conformational change. Overall, these findings showed that SMM **R3** can bind to both the GTP-bound and -unbound Rab10, thus meriting further structural optimization and experimental validation as a possible Rab10 inhibitor.

### Structural Analysis of Rab10 Interactions with SMMs

In an overview of the Rab10 protein (**Figure 1B** light blue), the binding site was illustrated in 3D (gray, CPK representation). The Switch I region (amino acids 31–43) is highlighted in green, the Switch II region (amino acids 66–82) in black, and the inter switch region (amino acids 44– 63) in magenta (PDB ID: 5SZJ). **Figure 1C-G** showed a detailed 3D view of the Rab10 GTP-binding site that highlights the interactions of active site residues with SMM **R1, R2, R3, R4** and **R5**, respectively. The 2D interactions for the top five compounds **R1-R5** for Rab10 were displayed in **Supplementary Figure S1**. As an example, compound **R1** fits well within the GTP-binding site, with favorable interactions including a crucial hydrogen bond with Thr23, essential for GTP-binding efficiency (**Figure 1**)^31^. Dynamic simulations for compound **R1-R5** showed high stability with an RMSD range of 2.3 Å to 3.2 Å (**Figure 1H-L**).

### ADME Properties and Toxicity Analysis

The ADME/Tox analyses using Schrödinger QikProp and SwissADME revealed that the top five compounds **R1-R5** (**Supplementary Table S2 and S3**) predicted good lipophilicity and water solubility, high gastrointestinal absorption, and compliance with Lipinski’s rule. The top-hit compounds **R1-R5** were further screened using the ProTox-II server to evaluate their potential toxicity. ProTox-II can predict the potential risks of carcinogenicity, immunotoxicity, mutagenicity, cytotoxicity, and hepatotoxicity, as well as estimate the LD50 value in mg/kg (body weight). The LD50 values categorize compounds into toxicity classes ranging from I to VI, where class I represents highly fatal compounds, and class VI indicates substances with minimal harmful risk upon ingestion^32^.

**R1** had an LD50 value of 1000 mg/kg, **R2** showed 500 mg/kg, **R3** exhibited 1190 mg/kg, **R4** displayed 722 mg/kg, and **R5** had an LD50 of 3000 mg/kg (**Supplementary Table S4**). All five compounds were classified under either toxicity class IV or class V, indicating moderate toxicity. The predicted LD50 values suggest favorable safety profiles, as these compounds were found to be inactive for most toxic endpoints, including hepatotoxicity, cardiotoxicity, immunotoxicity, cytotoxicity, and mutagenicity.

### Comparative Docking Predicts Preferential Interactions with GTP-bound Rab10

To assess predicted Rab-family and nucleotide-state preferences, compounds **R1-R5** were docked against Rab5, Rab8, Rab10, and Rab11 in GTP-bound (**Table 1**), GDP-bound (**Supplementary Table S5**), and nucleotide-free states (**Table 2**). Rab8 was selected as a closely related Rab10 family member to examine potential discrimination between related proteins^33^. Rab11 and Rab5 were included as representative regulators of recycling and early endosomal trafficking, extending the comparison to functionally relevant Rab proteins^34–36^.

**Table 1.**
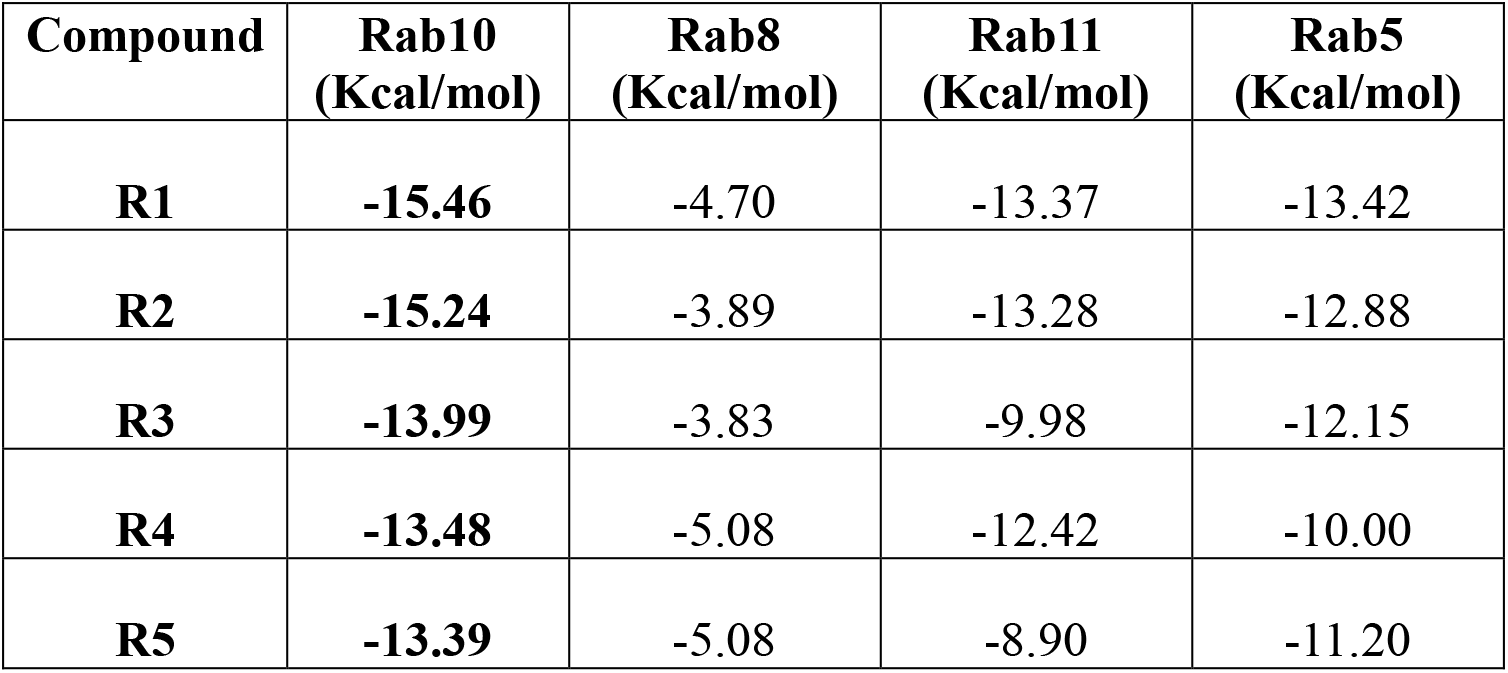
Predicted binding of R1-R5 to GTP-bound Rab10 and representative Rab-family proteins. Molecular docking scores (Kcal/mol) of compounds **R1-R5** against GTP-bound Rab10 (PDB: 5SZJ), Rab5 (PDB: 1RRP), Rab8 (PDB: 4LHZ) and Rab11 (PDB: 2D7C). Docking was performed using the same computational workflow with GTP retained in the nucleotide-binding site. More negative docking scores indicate more favorable predicted protein-ligand interactions.

| **Compound** | **Rab10<br>(Kcal/mol)** | **Rab8<br>(Kcal/mol)** | **Rab11<br>(Kcal/mol)** | **Rab5<br>(Kcal/mol)** |
|---|---|---|---|---|
| **R1** | **-15.46** | -4.70 | -13.37 | -13.42 |
| **R2** | **-15.24** | -3.89 | -13.28 | -12.88 |
| **R3** | **-13.99** | -3.83 | -9.98 | -12.15 |
| **R4** | **-13.48** | -5.08 | -12.42 | -10.00 |
| **R5** | **-13.39** | -5.08 | -8.90 | -11.20 |

**Table 2.** Predicted binding of R1-R5 to nucleotide free Rab10 and representative Rab-family proteins. Molecular docking scores (Kcal/mol) of compounds **R1-R5** against GTP-bound Rab10 (PDB: 5SZJ), Rab5 (PDB: 1RRP), Rab8 (PDB: 4LHZ) and Rab11 (PDB: 2D7C). Docking was performed using the same computational workflow with nucleotide free binding site. More negative docking scores indicate more favorable predicted protein– ligand interactions.

| **Compound** | **Rab10 (Kcal/mol)** | **Rab8 (Kcal/mol)** | **Rab11 (Kcal/mol)** | **Rab5 (Kcal/mol)** |
|---|---|---|---|---|
| **R1** | **-11.48** | -5.94 | -9.84 | -4.54 |
| **R2** | **-11.31** | -3.13 | -12.09 | -10.39 |
| **R3** | **-7.81** | -3.35 | -7.42 | -7.27 |
| **R4** | **-9.73** | -3.91 | -3.61 | -7.74 |
| **R5** | **-8.68** | -6.48 | -6.49 | -8.75 |

All five compounds scored most favorably against Rab10 within the GTP-bound panel, with scores ranging from −15.46 to −13.39 Kcal/mol. **R1** and **R2** yielded the most favorable Rab10 scores, at −15.46 and −15.24 Kcal/mol, respectively. Scores favored Rab10 over Rab8 by 8.31–11.35 Kcal/mol, over Rab11 by 1.06–4.49 Kcal/mol, and over Rab5 by 1.84–3.48 Kcal/mol. **R3** and **R5** showed the largest separation from Rab11, whereas **R4** displayed the smallest Rab10– Rab11 difference.

All compounds also scored more favorably against GTP-bound Rab10 than against its GDP-bound or nucleotide-free structures. **R3** exhibited the largest nucleotide-state differences, with scores of −13.99, −8.10, and −7.81 Kcal/mol for GTP-bound, GDP-bound, and nucleotide-free Rab10, respectively. In contrast, **R4** showed a smaller GTP–GDP difference of 0.96 Kcal/mol. Rab-family preferences were less consistent in the other states: **R1** and **R3** favored GDP-bound Rab5 over Rab10, **R2** favored nucleotide-free Rab11, and **R5** showed nearly equivalent scores for nucleotide-free Rab10 and Rab5. Collectively, these results support a predicted preference for GTP-bound Rab10 within the tested panel, with compound-dependent differences in Rab-family and nucleotide-state discrimination.

### The Binding Kinetics of Rab10 SMMs

SPR experiments were performed to directly assess the binding of **R1**-**R4** to Rab10 and Rab5 in the absence or presence of GTP. Without GTP, the compounds bound Rab10 with micromolar KD values: 37.1 µM for **R1**, 86 µM for **R2**, 46.9 µM for **R3**, and 70 µM (**Figure 2**) for **R4** and steady state profiles are included in the **Supplementary Figure S2**. The addition of GTP substantially enhanced the apparent affinity of every compound for Rab10. The KD values decreased to 12 nM for **R1**, 40 nM for **R2**, 0.7 nM for **R3**, and 2 nM for **R4**. ZCL278, used as a reference compound, showed weak millimolar binding to Rab10 both without GTP (10.57 mM) and with GTP (12.4 mM) (**Table 3**).

**Figure 2.**
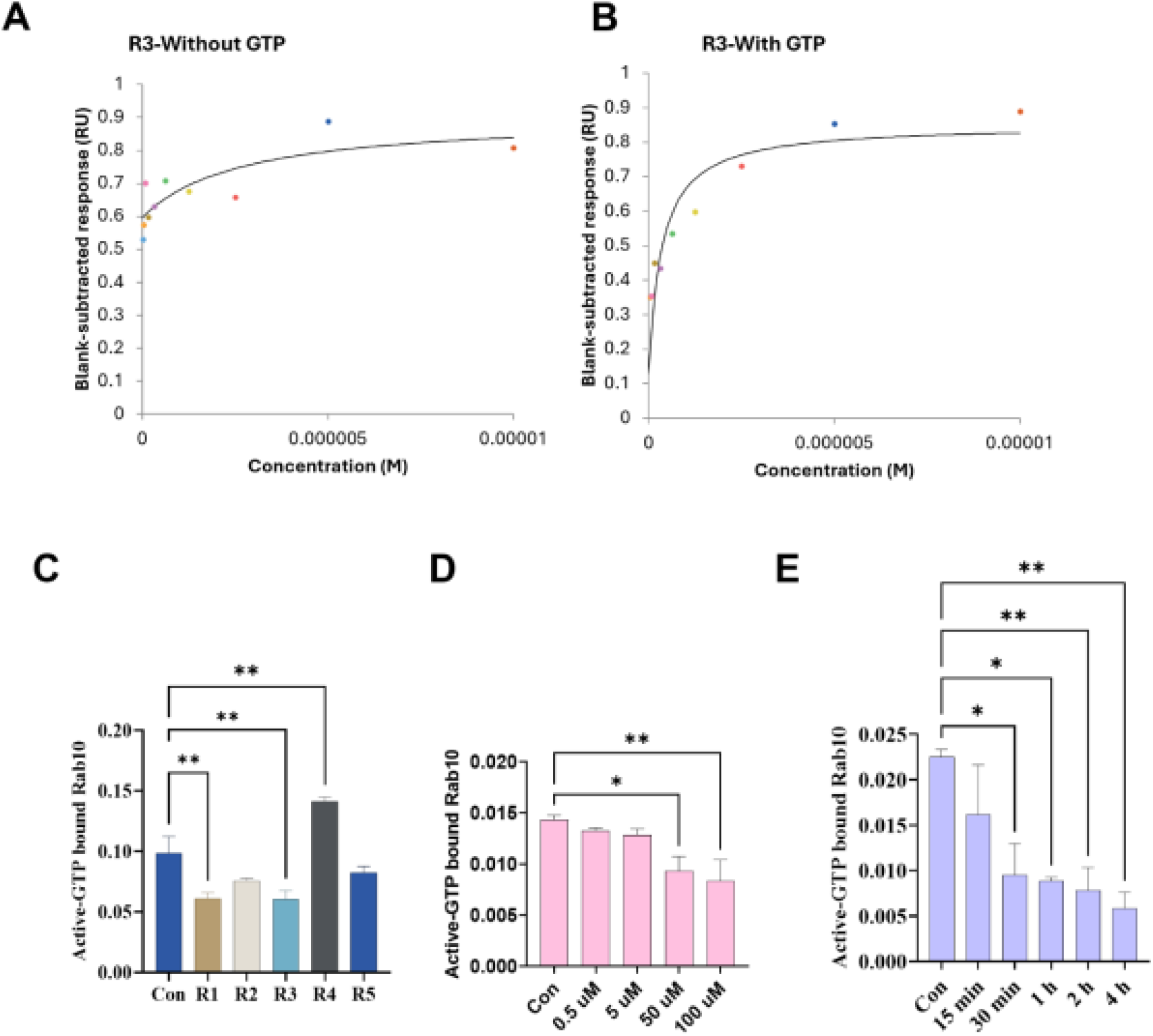
Biochemical validation of SMMs R1-R5 binding to Rab10 and their modulation of Rab10 activity. **(A-B)** SPR assessment of Rab10 binding affinities of compounds **R3** with and without GTP. With and with out GTP state state affinity of **R3** with Rab10. (**C**) Cell-based validation of **R1-R5** modulation of Rab10 activity by G-LISA measurement of GTP-bound levels in Rab10. **R1**, **R2**, **R3** and **R5** (at the concentration of 50 µM) inhibited Rab10 activation whereas **R4** promoted Rab10 activation. Control included untreated cellular lysates. **(D-E)** Dose- and Time-Dependent Inhibition of Rab10 Activity by **R3** in HEK293 Cells. **(D)** HEK293 cells were incubated for 2 h with varying concentrations of compound **R3** (0.5, 5, 50, and 100 μM) and GTP-bound Rab10 was measured by G-LISA. **(E**) In a time-dependent study, HEK293 cells were treated with 50 μM of **R3** for different durations (15, 30, 60, 120, and 240 min) and GTP-bound Rab10 was measured by G-LISA. All data are presented as mean±SEM from duplicates in at least two independent experiments. ANOVA compared treatments to their respective control p < 0.05 (*), and p < 0.0005 (***).

**Table 3.** Steady-state binding affinities of test compounds for Rab10 and Rab5 in the presence or absence of GTP. Equilibrium dissociation constants (KD) were determined by SPR using Rab10 or Rab5 in running buffer with or without GTP. **R1-R4** showed markedly enhanced binding to Rab10 in the presence of GTP, whereas their affinities for Rab5 were generally reduced or only modestly altered. **ZCL278** was included as a reference compound.

|  | **Rab10** |  | **Rab5** |  |
|---|---|---|---|---|
|  | **Without GTP** | **With GTP** | **Without GTP** | **With GTP** |
| **Compound** | **Steady state affinity (KD)** | **Steady state affinity (KD)** | **Steady state affinity (KD)** | **Steady state affinity (KD)** |
| **R1** | 37.1 µM | 12 nM | 18.4 µM | 492 µM |
| **R2** | 86 µM | 40 nM | 373 µM | 55.58 µM |
| **R3** | 46.9 µM | 0.7 nM | 36.9 µM | 201 µM |
| **R4** | 70 µM | 2 nM | 53.15 µM | 2.215 mM |
| **ZCL278** | 10.57 mM | 12.4 mM | 1.35 mM | 39.8 mM |

In contrast, GTP did not produce a similar overall enhancement in Rab5 binding. **R1**, **R3**, and **R4** showed weaker apparent affinity for Rab5 in the presence of GTP, while **R2** showed improved Rab5 binding. Under GTP-containing conditions, **R1-R4** showed substantially stronger apparent affinities for Rab10 than for Rab5, with **R3** and **R4** exhibiting the greatest difference between Rab10 and Rab5 binding (**Table 3**).

### Biochemical Targets of Rab10 SMMs in HEK293 Cells

To determine the biochemical targets of **R1-R5** in modulating Rab10 activities in HEK293 cells, we developed a Rab10 ***<u>C</u>***ell-based ***<u>G</u>***TP ***<u>Q</u>***uantification ***<u>E</u>***nzyme-***l***inked ***i***mmuno***<u>s</u>***orbent ***<u>a</u>***ssay (Rab10-CGQ-ELISA). This analytic biochemical methodology allowed us to systematically assess the effects of **R1-R5** on Rab10 activation in HEK293 cells. At the basal state, **R1, R2, R3, and R5** suppressed Rab10 activation, shown as reduced GTP-bound Rab10, whereas **R4** enhanced activation (**Figure 2C**). These observations indicated that distinct structural features among these compounds may underlie their opposing pharmacological action modes.

To further characterize SMM **R3**, we evaluated its dose- and time-dependent effects on Rab10 activation. A clear dose-dependent response was observed, where treatment from low (0.5 µM) to high concentrations (100 µM) led to progressively reduced GTP-bound Rab10 (**Figure 2D**). Similarly, temporal analysis revealed a reduction of Rab10 activation between 15 minutes and 4 h of treatment (**Figure 2E**). These findings highlight the dynamic nature of **R3**’s interaction with Rab10.

### Rab10 SMMs Modify Early Endosome Organization

Rab10 regulates membrane trafficking within cells, directing vesicle movements between compartments, especially the early and recycling endosomes^35^. Therefore, we sought to determine endosome organization when HEK293 cells were treated with Rab10 SMMs at a concentration of 50 µM for 2 h (**Figure 3A-C**).

**Figure 3.**
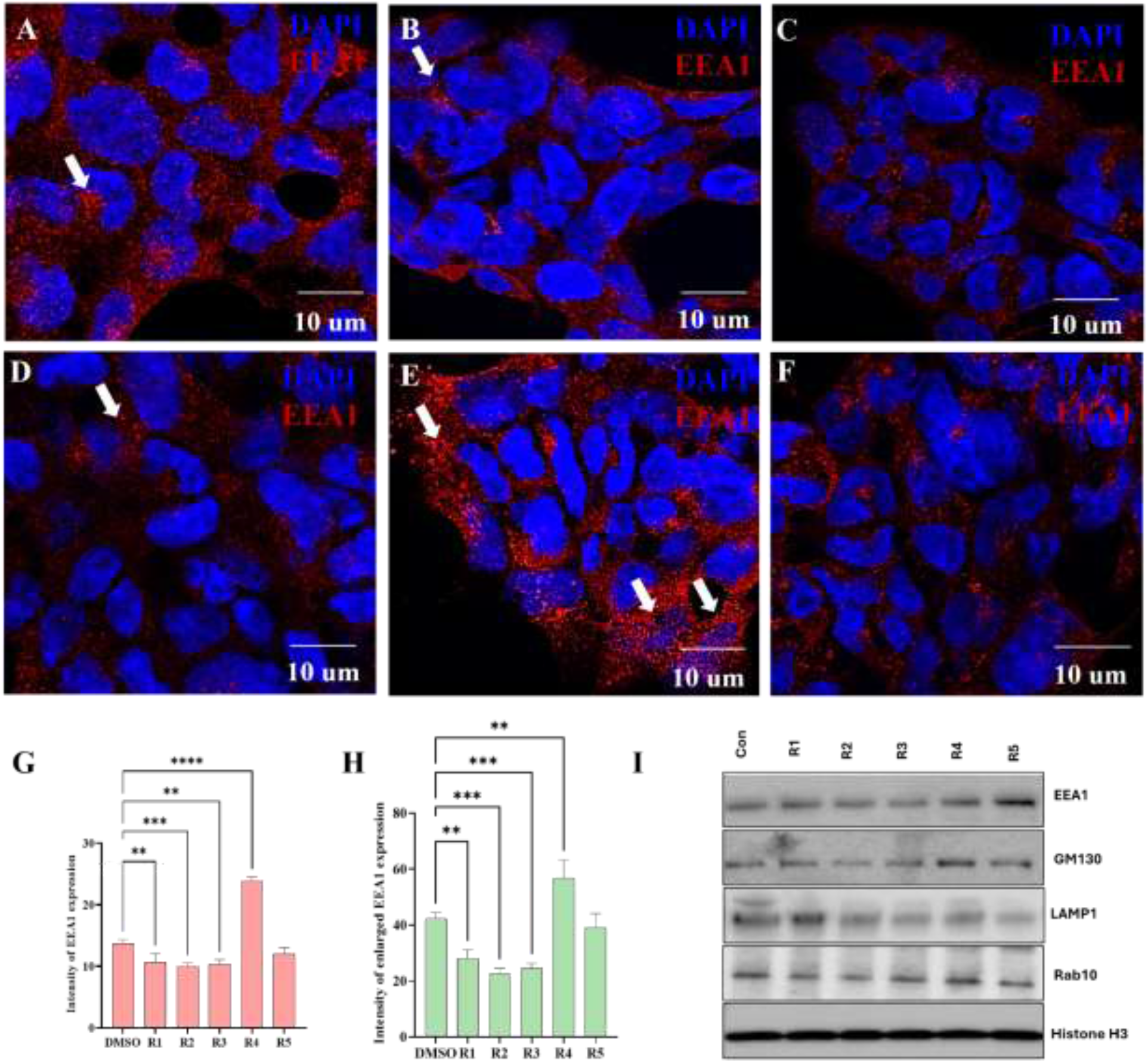
Effects of Rab10 SMMs on endolysosomal homeostasis in HEK293 cells. Treatment with compounds **R1-R3** and **R5** at **50 µM (A-C and F)** produced obvious changes in EEA1-positive endosomal morphology compared with the control. In contrast, treatment with **R4 (D)** was associated with enlarged EEA1-positive endosomal structures (white arrows), suggesting an effect on endosomal organization. **(G)** Quantification of EEA1 signal in HEK293 cells treated with **R1-R5. (H)** Quantification of the enlarged endosomal effects of **R1-R5** in HEK293 cells treated. Data are presented as mean ± SD, with statistical significance defined as p < 0.005 (**), p < 0.0005 (***), and p < 0.0001 (****). Scale bar = 10 µm. **(I)** Western blot analysis of EEA1, GM130, LAMP1 and total Rab10 in HEK293 cells. HEK293 cells were treated with 50 µM of SMMs **R1-R5.** Histone-H3 was used as a loading control.

**Figure 3A** showed the effects of **R3,** as a representative Rab10 inhibitor, and **R4**, a representative Rab10 activator, on the endosomal organization. Compared to control, cells treated with **R3** showed a reduced number of endosomes as determined by anti-EEA1 immunostaining (**Figure 3A, B and G**; Arrows: enlarged endosomes). **R1-R3** and **R5** showed similar inhibitory effects to **R3** on Rab10-mediated endosome organizations.

On the other hand, **R4** treatment led to a significantly high number of endosomes, as well as the enlarged endosomes, suggesting that **R4** may activate Rab10 and shift the homeostatic progression of endosomes to sort and traffic cargo (**Figure 3C**; Arrows: enlarged endosomes). Quantification of the EEA1 expression in HEK293 cells further validated the observed differences in endosomal organization across the five top ranking **R1-R5** SMMs.

The different effects of **R1-R5** SMMs on endolysosomal network (ELN) proteins, such as EEA1 and LAMP1 expressions (**Supplementary Figure S3**), were additionally evaluated by Western blots. **R1-R5**, except for **R4**, reduced EEA1 expression (**Figure 3I**). A very similar trend was observed for LAMP1 and quantification data included in the **Supplementary Figure S4.**

### APP As a Target of Rab10 SMMs

In Chinese Hamster Ovary (CHO) cells overexpressing APP and presenilin-1 (PS70), APP undergoes a maturation process in the Golgi apparatus, including glycosylation^37^. Consistent with the literature, overexpressed APP was enriched in the perinuclear compartment of the Golgi apparatus (**Figure 4A: Arrows**). Treatment of CHO cells with brefeldin A (BFA) interrupted Golgi organization and redistributed APP to ER-like structures (**Figure 4B**: Asterisks). **R3** treatment appeared to promote intracellular trafficking, as APP was not as concentrated in Golgi as in untreated PS70 cells but also not enriched in ER as in BFA-treated cells. Rather, APP in **R3-** treated PS70 cells appeared to be distributed to the ER, Golgi, and ELN-like compartments, respectively (**Figure 4C**). Collectively, compared to untreated and BFA treated cells, **R3** reduced the total APP expression as revealed by anti-APP (CT695) immunoreactivity (**Figure 4G**).

**Figure 4.**
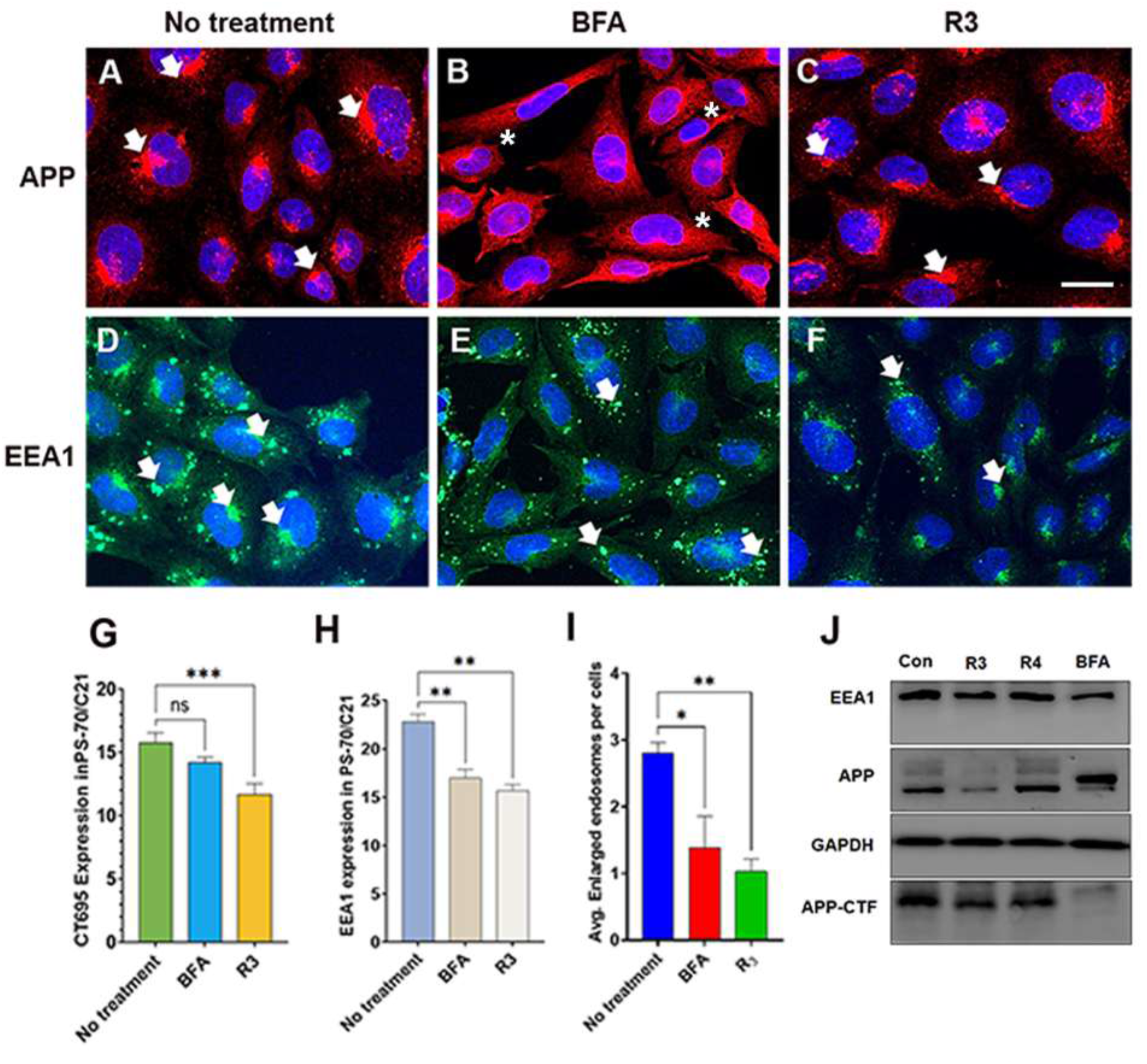
Effects of Rab10 SMM R3 on APP processing and endolysosomal homeostasis in CHO cells overexpressing APP and presenilin-1. (**A-C**) Effects of **R3** on APP distribution in CHO cells overexpressing APP and presenilin-1. (**A**) Control; **(B)** Brefeldin A (BFA) treatment, and **(C) R3** treatment. Nuclei were counterstained with DAPI (blue). White arrows indicate representative CT695-positive APP puncta or intracellular accumulations. In control cells with no **R3** or BFA treatments, overexpressed APP displayed prominent perinuclear clustering of aggregation. BFA caused a dramatical redistribution of CT695-positive APP to ER-like organization (Asterisks). **R3** decentralized CT695-positive APP towards endolysosomal vesicles. (**D-F)** Effects of **R3** on EEA1 distribution in CHO cells overexpressing APP and presenilin-1. **(D)** Control; **(E)** Brefeldin A (BFA) treatment, and **(F) R3** treatment. Nuclei were counterstained with DAPI (blue). In control with no treatment of BFA or **R3,** numerous EEA1-positive vesicular structures were detected throughout the cytoplasm (white arrows). Treatment with BFA or **R3** resulted in changes in EEA1-positive vesicle distribution and abundance. White arrows highlight EEA1-positive vesicular structures. (**G**) Quantification of CT695 fluorescence intensity relative to untreated control cells. Compared with untreated and BFA treated cells, **R3** treatments reduced intracellular CT695 accumulation. (**H**) Quantification of EEA1 fluorescence intensity. Treatment with BFA or R3 decreased EEA1-positive endosomal signal relative to untreated cells. (**I**) Quantification of EEA1 fluorescence intensity of enlarged endosomes. Treatment with BFA or **R3** decreased EEA1-positive enlarged endosomal signal relative to untreated cells. (**J**) Western blot analysis of CT695-positive APP and EEA1 in untreated, BFA-treated, and **R3-**treated cells. BFA altered APP maturation whereas **R3** reduced APP expression. Data are presented as mean ± SEM from at least three independent experiments. GAPDH was used as loading control. Statistical significance was determined using one-way ANOVA followed by an appropriate post hoc test (\**P* < 0.05, \*\**P* < 0.01, \*\*\**P* < 0.001 versus untreated control). Scale bar, 10 μm.

Supporting the targeting of **R3** on Rab10 in the ELN system, we further assessed the EEA1 distribution in PS70 cells. As expected, the ELN system was likely overwhelmed by the overexpression of APP and presenilin-1 that EEA1 showed many enlarged endosome-like structures (**Figure 4D**; Arrows: enlarged endosomes). Both **R3** and BFA treatment reduced the total number of endosomes as well as enlarged endosome-like structures (**Figure 4E and F**; Arrows: enlarged endosomes**; See also Figure 4H and I** quantification of the data).

Corroborating the effects of Rab10 SMMs on intracellular trafficking systems that carry protein cargo such as APP, **R3** reduced full-length APP expression as well as APP carboxyl terminal fragments (CTFs) as shown (**Figure 4J**). Here, we observed the clear functional separation of Rab10 SMMs such as **R3** from Golgi disrupter BFA. Treatment of CHO cells with BFA dramatically changed the APP maturation pattern and the inhibition of glycosylation. However, **R3** or **R4** did not alter the APP maturation pattern, indicating that their primary targets are not Golgi structures (**Figure 4J**). Moreover, while BFA prohibited the APP CTF formation, likely due to the total inhibition of Golgi function, **R3** or **R4** decreased the secretase-mediated cleavage, resulting in the reduction of APP CTF (**Figure 4J**).

### Neuroprotective Effects of Rab10 SMMs

PC12 cells acquire a neuron-like phenotype upon sustained nerve growth factor (NGF) stimulation and undergo apoptosis following NGF withdrawal. Previous studies have shown that NGF withdrawal-induced neurodegeneration is accompanied by altered APP processing and can be attenuated by inhibiting amyloidogenesis^38^. Given Rab10’s involvement in APP processing and intracellular trafficking, we investigated whether Rab10 inhibition could mitigate neuronal degeneration following NGF withdrawal. PC12 cells were stained with phalloidin to visualize F-actin and counterstained with DAPI to assess nuclear morphology (**Figure 5A-E**). Under continuous NGF exposure, cells maintained intact nuclei and a differentiated morphology with prominent neurite extensions (**Figure 5A**, arrows). In contrast, NGF withdrawal reduced cell density and disrupted F-actin organization, with some cells exhibiting no detectable F-actin staining (**Figure 5B**, double asterisks). Cell shrinkage and diminished cytoskeletal staining were consistent with cytoskeletal deterioration and loss of cell viability, supporting the requirement for NGF in maintaining the survival and morphology of differentiated PC12 cells.

**Figure 5.**
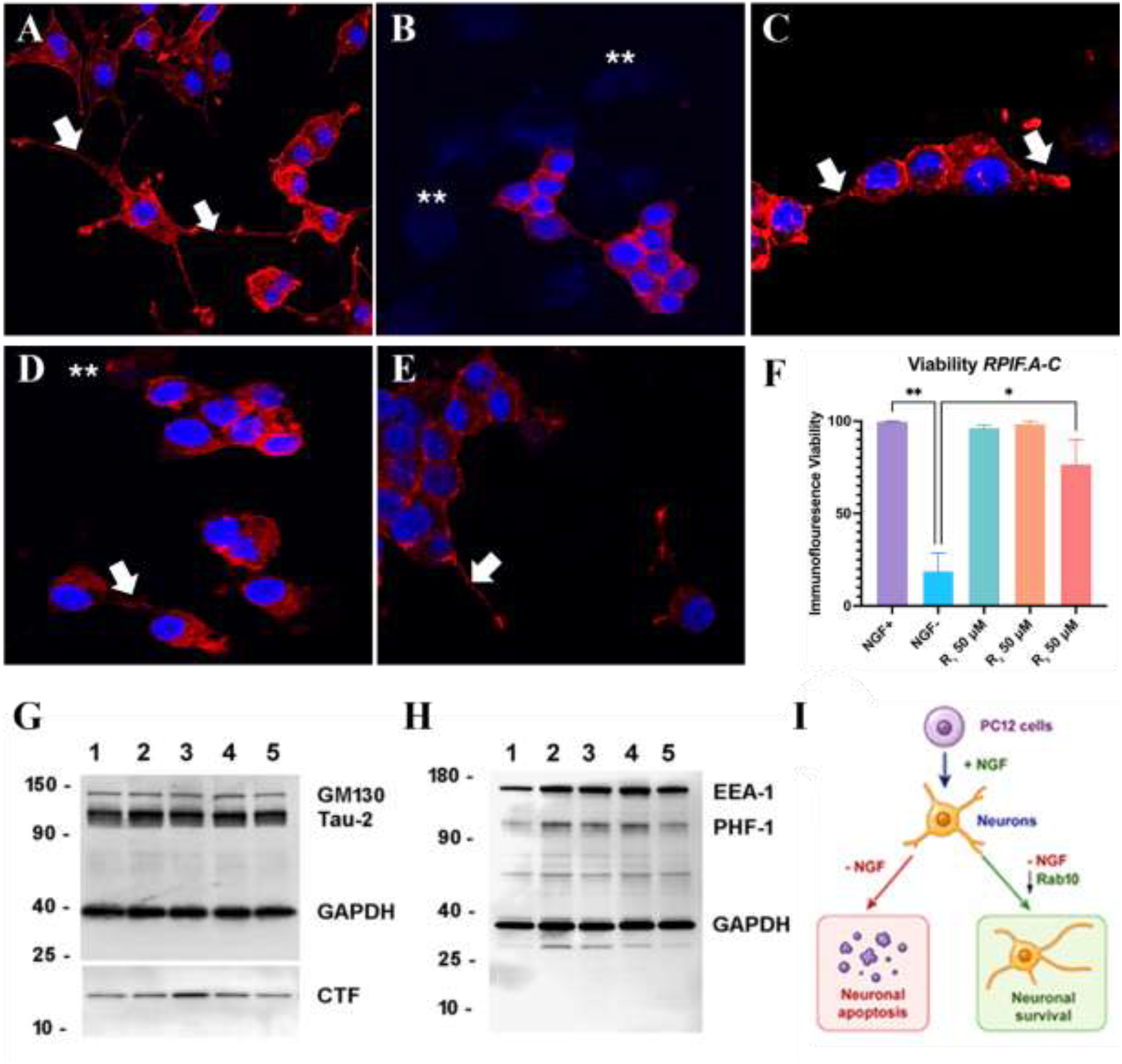
Rab10 modulation reduces nerve growth factor (NGF) withdrawal-induced neuronal degeneration in PC12 cells. **(A-E)** Representative immunofluorescence images of NGF-differentiated PC12 cells maintained with NGF **(A)** subjected to NGF withdrawal (B), or treated with 50 μM **R1 (C), R2 (D), or R3 (E)** following NGF withdrawal. F-actin is shown in red, and nuclei are stained with DAPI (blue). Arrowheads indicate stubby protrusions; arrows, neurite extensions; asterisks, cell shrinkage; and double asterisks, cells lacking detectable F-actin staining. **(F)** Quantification of cell survival under the indicated treatment conditions, normalized to the untreated control group. **(G)** Western blot analysis of APP C-terminal fragments (APP-CTFs), total tau detected using the Tau-2 antibody, and GM130. **(H)** Western blot analysis of phosphorylated tau detected using the PHF-1 antibody (pSer396/pSer404) and EEA1. For **(G)** and **(H),** lanes correspond to undifferentiated PC12 cells (lane 1), NGF-differentiated PC12 cells maintained with NGF (lane 2), cells subjected to NGF withdrawal (lane 3), and cells treated with 10 μM (lane 4) or 50 μM (lane 5) **R3** following NGF withdrawal. GAPDH served as the loading control. **(I)** Schematic illustrating NGF withdrawal-induced neurodegeneration and the proposed protective effects of Rab10 inhibition on neuronal survival. Data are presented as mean ± SEM from multiple independent experiments. *P* < 0.05 for the indicated comparisons.

Treatment with the Rab10 SMMs **R1-R3** showed neuroprotective effects following NGF withdrawal. Compared with **R2**, **R1** and **R3** produced greater improvements in cell morphology, including increased cell density, improved F-actin organization, and partial recovery of neurite extensions (**Figure 5C-E**). Consistent with these observations, quantitative analysis showed that NGF withdrawal markedly reduced cell survival, whereas treatment with 50 μM **R3** significantly improved survival (**Figure 5F**).

We finally examined APP and tau protein expression and posttranslational modification associated with NGF withdrawal-induced apoptosis and the effects of Rab10 modulation. APP-CTF was increased in differentiated PC12 cells following NGF withdrawal, but this trend was reversed by **R3** (**Figure 5G**: Compare Lane 5 with Lane 3). Interestingly, phosphorylated tau was increased upon NGF stimulation which remained at an elevated level following NGF withdrawal (**Figure 5G**: Compare Lane 3 with Lane 2). However, **R3** treatment was able to reduce tau phosphorylation (**Figure 5H**: Compare Lane 5 with Lane 3). Overall, these findings indicate that Rab10 inhibition protects PC12 cells from NGF withdrawal-induced cytoskeletal disruption and cell death (**Figure 5I**) with the mechanisms potentially involving APP and tau modification.

## Discussion

Dysregulated endosomal trafficking contributes to AD related cellular dysfunction by influencing APP processing, intracellular cargo sorting, and neuronal homeostasis^3,7,13,19^. Human genetic studies implicating RAB10 in AD resilience provide the rationale for investigating its pharmacological modulation^15,16^. Here, we combined structure-based virtual screening, molecular docking, MD simulations, SPR, and cellular assays to identify candidate Rab10 SMMs.

Virtual screening identified five top candidates, **R1-R5,** with favorable predicted interactions near the Rab10 nucleotide-binding region. MD simulations supported the persistence of the modeled protein–ligand complexes. Computational ADME and toxicity assessments provided additional rationale for compound prioritization.

Comparative docking suggested that both Rab identity and nucleotide context influence predicted compound recognition. Rab8 was included as a closely related Rab protein, while Rab5 and Rab11 provided comparisons with regulators of early endosomal and recycling pathways^11,12,36^. All five compounds scored most favorably against Rab10 within the GTP-associated panel. The largest score differences occurred against Rab8, whereas smaller, compound-dependent differences were observed against Rab5 and Rab11. These findings suggest that related Rab proteins may differ in their capacity to accommodate the compounds within the modeled binding region.

SPR provided complementary evidence of GTP-associated enhancement of apparent binding. Without added GTP, **R1–R4** exhibited micromolar apparent dissociation constants, whereas GTP-containing conditions yielded nanomolar apparent values. Under the latter conditions, all four compounds showed stronger apparent binding to Rab10 than to Rab5. Together with the docking results, these observations support the hypothesis that a GTP-associated Rab10 environment favors compound recognition.

Cellular assays reveal distinct functional profiles despite the compounds shared predicted preference for GTP-associated Rab10. **R1, R2, R3,** and **R5** reduced GTP-bound Rab10, whereas **R4** increased it. The opposing effect of **R4** further illustrates why docking scores and binding affinity alone cannot establish functional mechanism.

The observed cellular phenotypes were consistent with perturbation of trafficking-associated processes. Previous work established a role for Rab10 in membrane transport through early endosomes in polarized epithelial cells^34^. In our HEK293 experiments, **R1-R3** and **R5** altered EEA1 abundance and EEA1-positive endosomal morphology, whereas **R4** was associated with enlarged endosomal structures. Changes in LAMP1 abundance additionally suggested effects on endolysosomal organization. These observations are relevant to the broader involvement of Rab-dependent trafficking in neurodegeneration^13,19^.

In APP/presenilin-1-expressing CHO cells, **R3** redistributed intracellular APP, reduced full-length APP and APP-CTFs, and decreased enlarged endosome-like structures. These findings are consistent with the connection between APP metabolism and endolysosomal dysfunction^7^. They also complement previous genetic evidence showing that RAB10 knockdown reduces Aβ42 production and the Aβ42/Aβ40 ratio, whereas overexpression increases the ratio and β-CTF abundance^15^. However, our experiments measured APP-related phenotypes rather than Aβ production and therefore do not establish reduced amyloidogenesis. The effects of **R3** differed from the pronounced APP redistribution and maturation changes associated with brefeldin A, suggesting that the compounds did not produce identical cellular responses.

**R3** also attenuated cytoskeletal deterioration and improved survival following NGF withdrawal in differentiated PC12 cells, accompanied by reduced APP-CTF accumulation and phosphorylated tau signals. Previous work linked NGF deprivation-induced PC12 cell death to activation of amyloidogenic APP processing^38^, providing a relevant context for these observations.

In conclusion, this study identified novel Rab10 SMMs that demonstrated preferential interactions with GTP-associated Rab10. Among the compounds, **R3** reduced cellular GTP-bound Rab10, altered endolysosomal organization and APP-related phenotypes, and attenuated NGF withdrawal-induced degeneration in differentiated PC12 cells. These findings support further development of **R3** as a candidate chemical biology probe for investigating Rab10-associated trafficking and AD-relevant cellular dysfunction.

## Statements and Declarations

### Author contributions

**Satyaveni Malasala:** Data curation and analysis, writing, and editing

**Nathan Cox:** Data curation and analysis of PC-12 cell treatments data and editing

**Grady Sellman:** Data curation and analysis of SPR and docking studies

**Qun Lu:** Conceptualization and study design, writing, and editing

**Yan-Hua Chen:** Conceptualization and study design, writing, and editing

## Acknowledgments

We thank Rodney Tatum and Christi Boykin for their technical assistance.

## Data and materials availability

The original contributions presented in the study are included in the article/supplementary material. Further inquiries can be directed to the corresponding author.

---

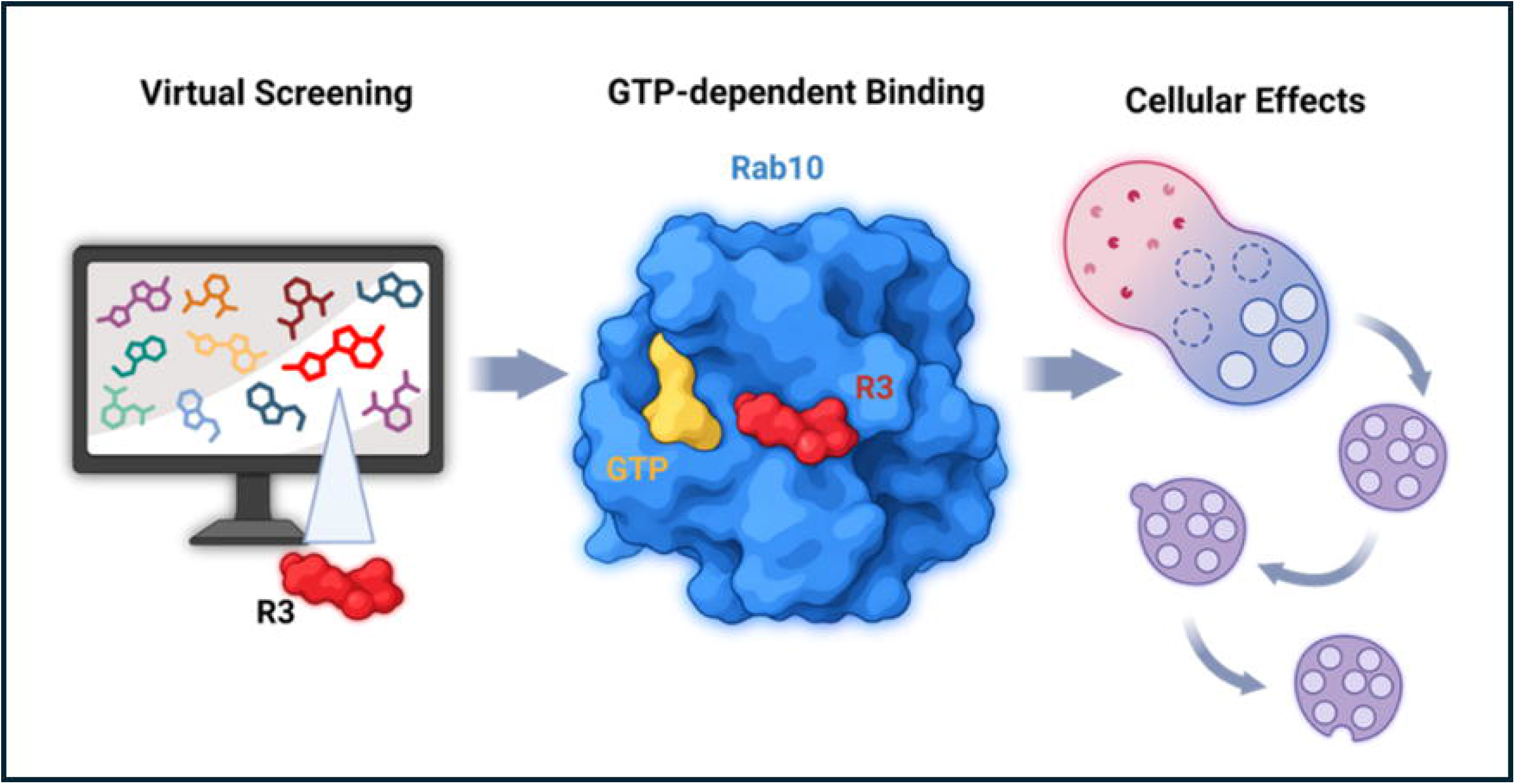

