## Supporting information for "Structure-Based Discovery of GTP-Enhanced Rab10 Modulators and Their Effects on Alzheimer’s Disease-Relevant Cellular Phenotypes"

### Supplementary Figures and Tables

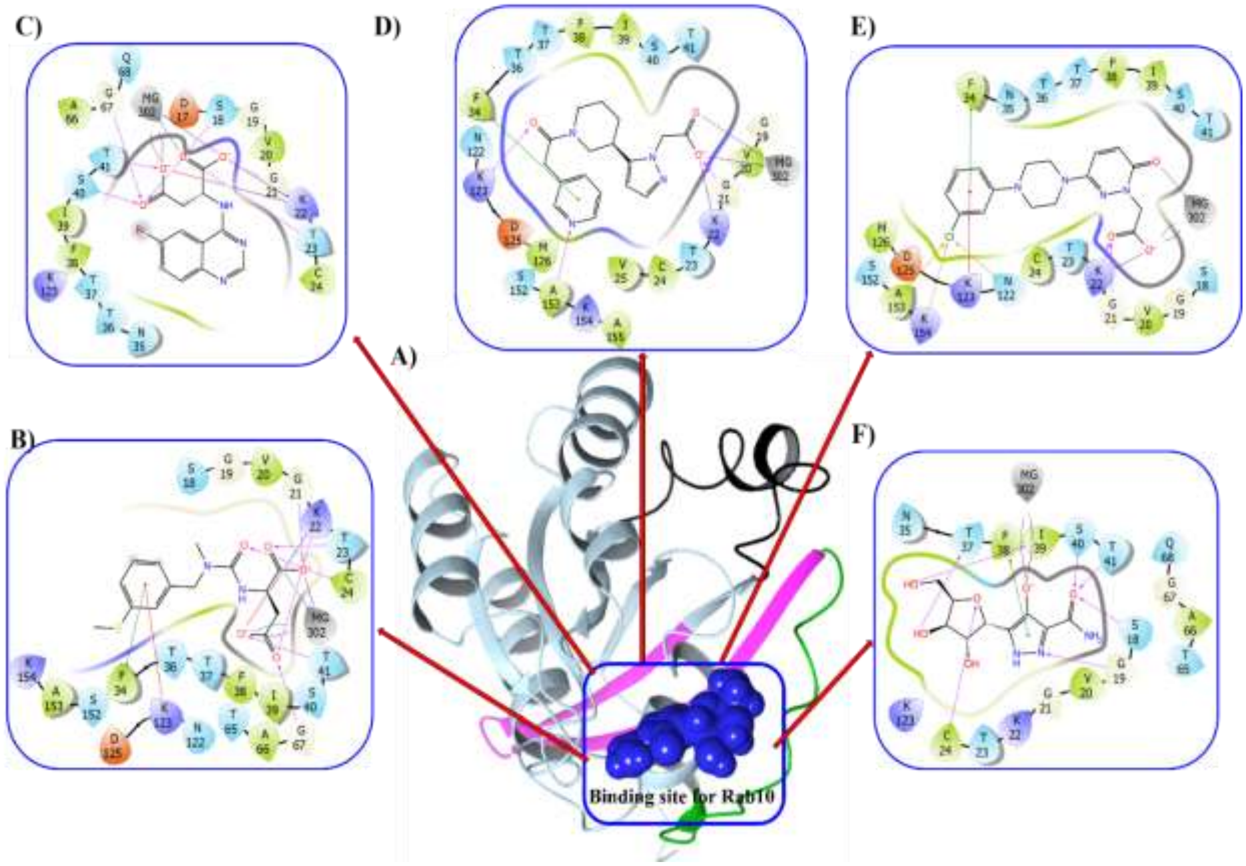


**Supplementary Figure S1. Superimpose of top five compounds R1-R5 and their interactions with Rab10. (A)** A 3D structural illustration of Rab10 is shown in transparent cyan, with specific regions highlighted: switch 1 (residues 31-43) in green, switch 2 (residues 66-82) in black, and the interswitch region (residues 44-63) in pink (PDB: 5SZJ). The predicted GTP binding site is represented in blue using the CPK model. **(B-E)** 2D representations of the active site binding residues of Rab10 interacting with **R1, R2, R3, R4,** and **R5** at the GTP site. Magenta arrows indicate hydrogen bonds. Binding site residues are color-coded based on interaction type: hydrophobic (green), polar (cyan), positively charged (violet), and negatively charged (red).

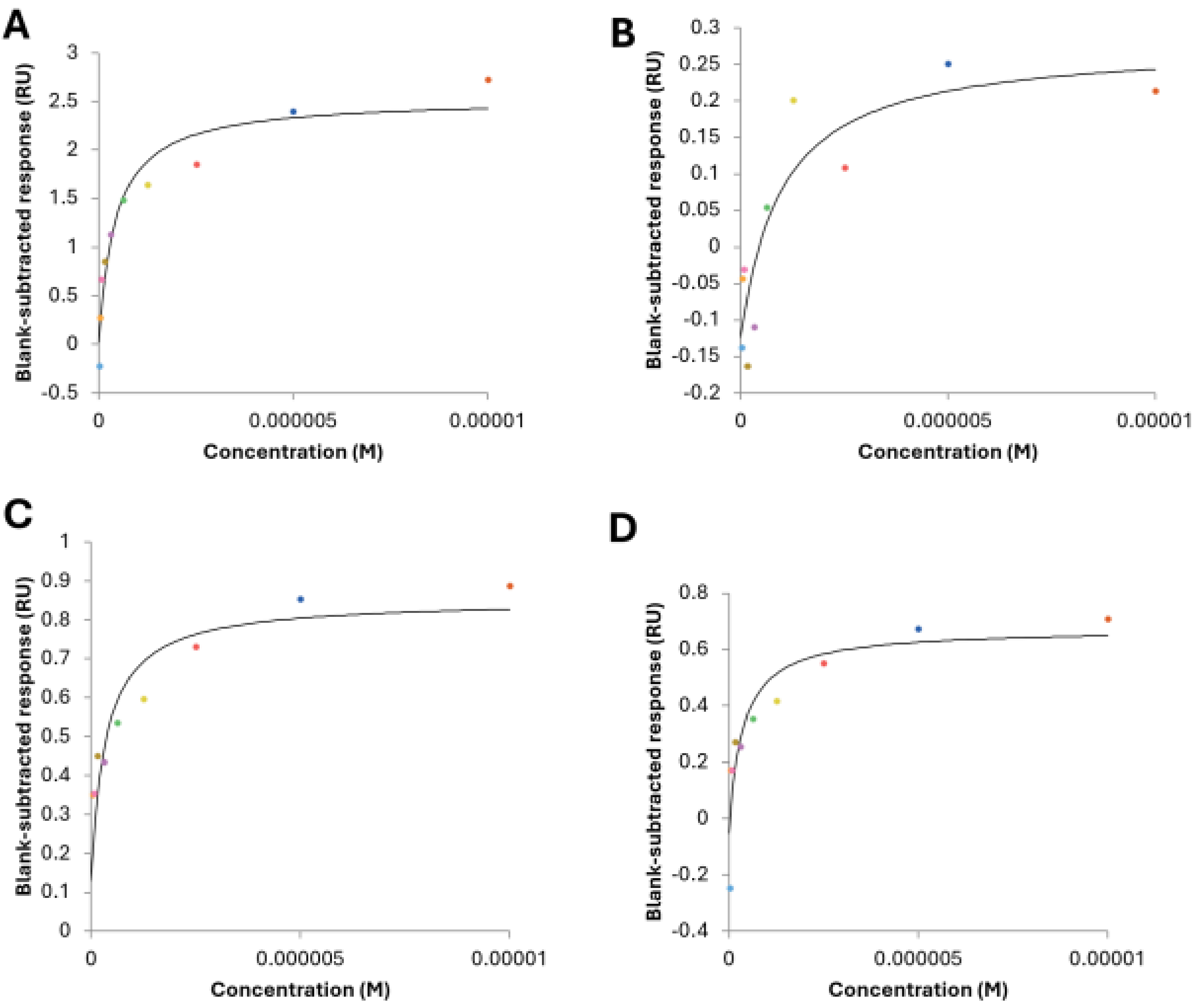


**Supplementary Figure S2 Steady-state SPR binding analysis of Rab10 small-molecule modulators in the presence of GTP (R1-R4).** Steady-state binding responses of compounds **R1-R4** to immobilized Rab10 were measured by surface plasmon resonance using running buffer supplemented with GTP. Colored symbols represent the steady-state, reference-subtracted SPR response at each analyte concentration, and solid lines represent nonlinear regression fits to a steady-state binding model.

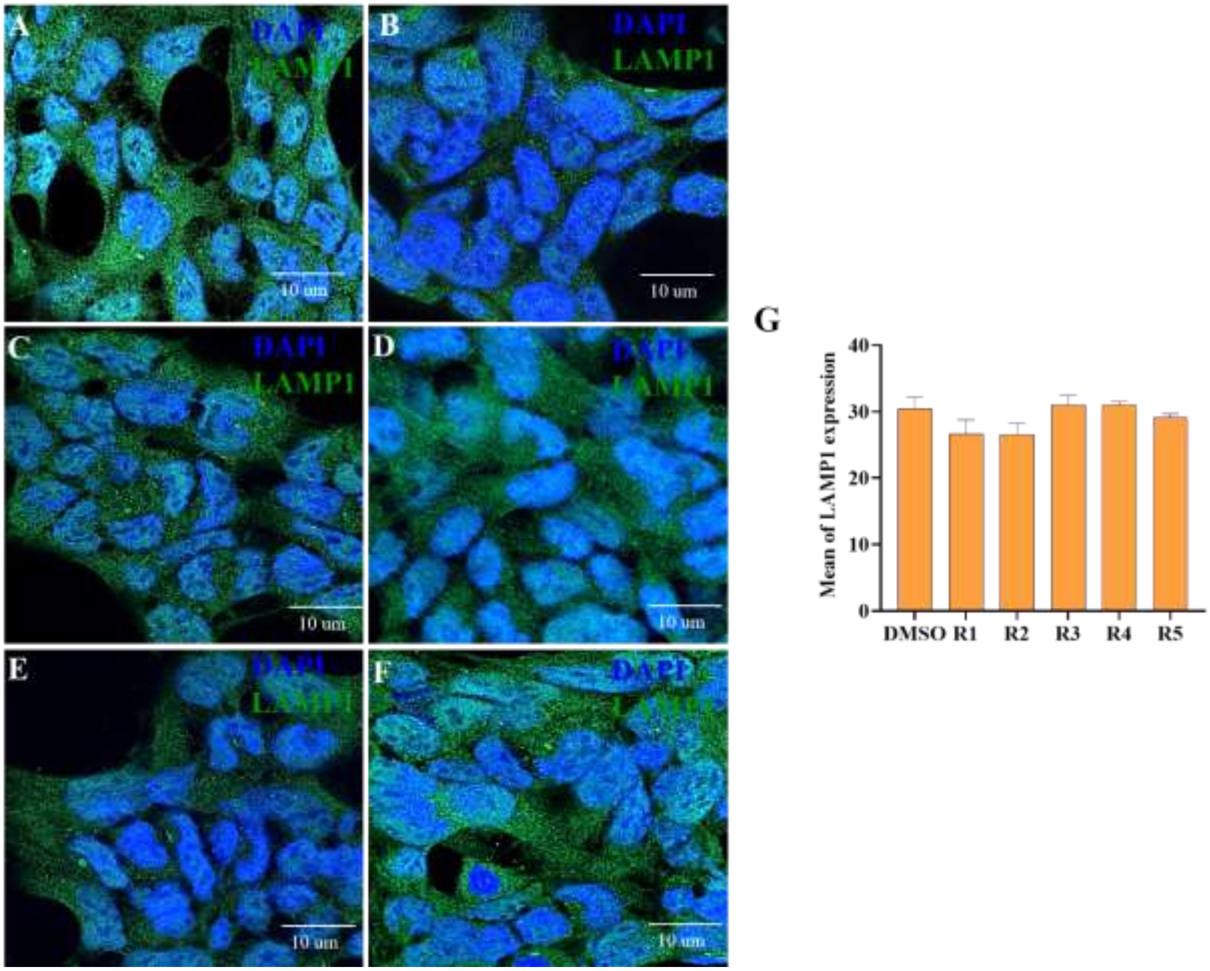


**Supplementary Figure S3. Late Endosomal Expression in HEK293 Cells.** In this study, we evaluated late endosomal expression in HEK293 cells following a 2-hour treatment with the top five identified compounds. After treatment, cells were fixed with 4% paraformaldehyde and immunostained for the late endosomal marker LAMP1. DMSO-treated cells served as the control **(A)**. Treatment with compounds **R1-R5** at a concentration of 50 µM **(B-F)** did not induce significant changes in late endosomal morphology or LAMP1 expression, indicating no observable toxicity or disruption to endosomal structure. Images were captured at 63× magnification using a Zeiss Axio Observer microscope. Panel **(G)** quantifies LAMP1 expression across the top four compounds, with DMSO as the control. Data are presented as mean ± SD and Scale bar = 10 µm.

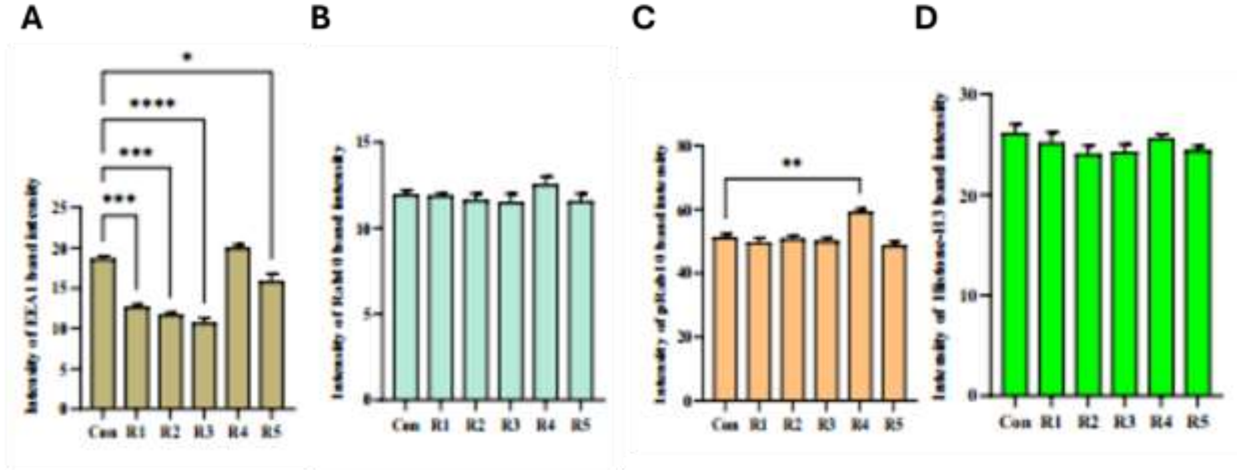


**Supplementary Figure S4. Western Blot data analysis quantification. A-D)** The expression levels of EEA1, LAMP1, Rab10, and Histone H3 were quantified, and the results further support the conclusions drawn from the study. Data are presented as mean ± SD, with statistical significance defined as $p < 0.003$ (**), $p < 0.0005$ (***), and $p < 0.0001$ (****).

**Supplementary Table S1. Identified top 100 small molecules targeting Rab10.**

| S. No | Molecule id | Docking Score | Structure | Molecular weight | Rule of Five Violation |
|---|---|---|---|---|---|
| 1. | ZINC000426465158 | -15.467 | 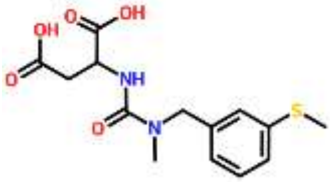 | 326.367 | 0 |
| 2. | ZINC000000140154 | -15.247 | 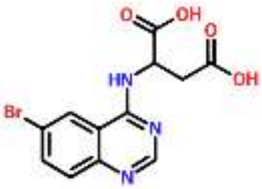 | 340.133 | 0 |
| 3. | ZINC000002132878 | -14.959 | 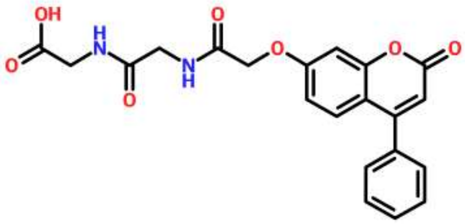 | 410.382 | 0 |

| | | | | | |
|---|---|---|---|---|---|
| 4. | ZINC000408592938 | -14.857 | 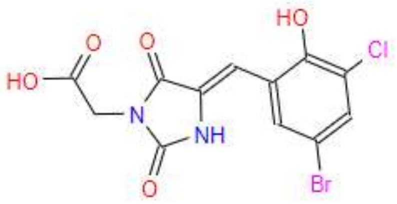 | 375.563 | 0 |
| 5. | | -14.351 | 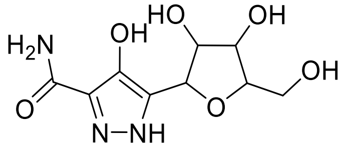 | 259.22 | 0 |
| 6. | ZINC000257273335 | -14.107 | 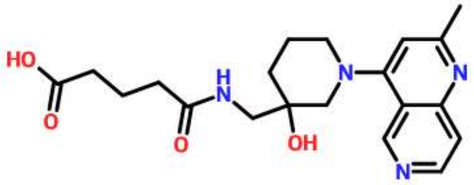 | 386.45 | 0 |
| 7. | ZINC000408590335 | -14.007 | 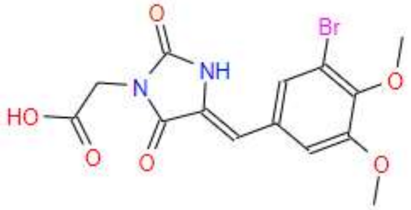 | 385.17 | 0 |
| 8. | ZINC000071404939 | -13.788 | 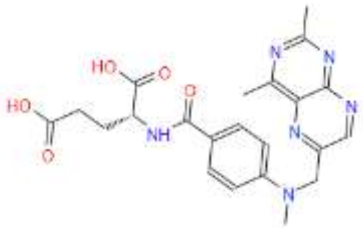 | 452.469 | 1 |
| 9. | ZINC000426612999 | -13.641 | 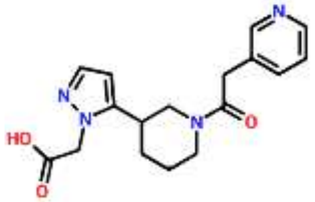 | 328.37 | 0 |
| 10. | ZINC000015857628 | -13.584 | 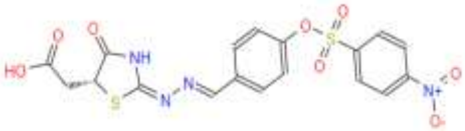 | 478.451 | 1 |
| 11. | ZINC000408521643 | -13.5 | 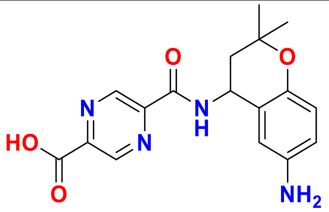 | 342.36 | 0 |

| | | | | | |
|---|---|---|---|---|---|
| 12. | ZINC000072220539 | -13.448 | 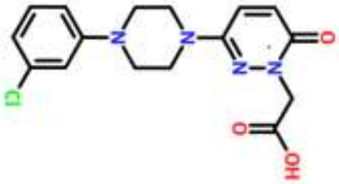 | 348.788 | 0 |
| 13. | ZINC000055090686 | -13.395 | 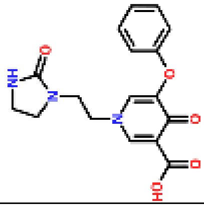 | 343.338 | 0 |
| 14. | ZINC000328577255 | -13.316 | 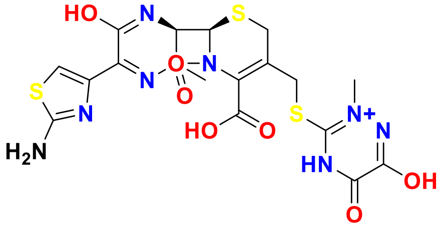 | 543.05 | 1 |
| 15. | ZINC000101273639 | -13.252 | 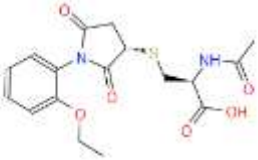 | 380.415 | 0 |
| 16. | ZINC000098212096 | -13.205 | 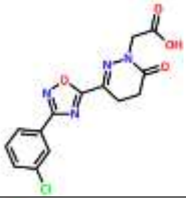 | 334.718 | 0 |
| 17. | ZINC000426373857 | -13.161 | 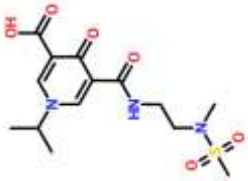 | 359.396 | 0 |
| 18. | ZINC000101273603 | -13.153 | 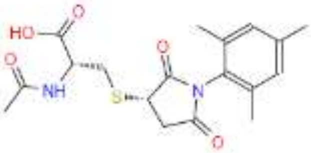 | 378.442 | 0 |
| 19. | ZINC000575414433 | -13.102 | 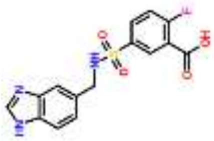 | 349.336 | 0 |

| | | | | | |
|---|---|---|---|---|---|
| 20. | ZINC000254713690 | -13.013 | 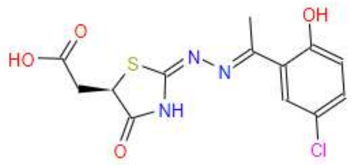 | 341.77 | 0 |
| 21. | ZINC000622000444 | -13.048 | 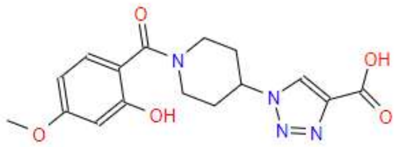 | 346.342 | 0 |
| 22. | ZINC000408592159 | -12.97 | 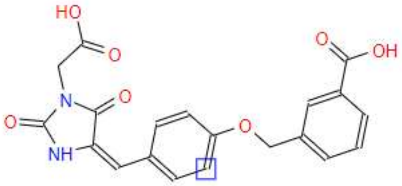 | 396.356 | 0 |
| 23. | ZINC000663103874 | -12.853 | 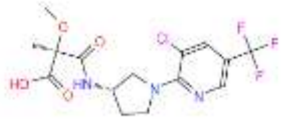 | 395.765 | 1 |
| 24. | ZINC000237446140 | -12.846 | 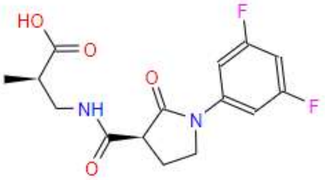 | 326.299 | 0 |
| 25. | ZINC000012884457 | -12.837 | 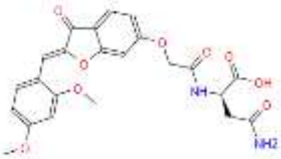 | 470.435 | 1 |
| 26. | ZINC000043701879 | -12.808 | 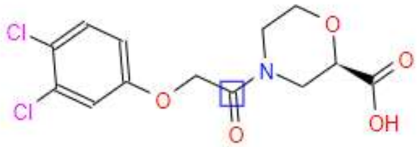 | 334.155 | 0 |
| 27. | ZINC000408593743 | -12.789 | 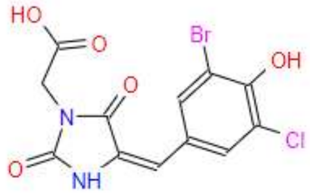 | 375.563 | 0 |
| 28. | ZINC000580097666 | -12.78 | 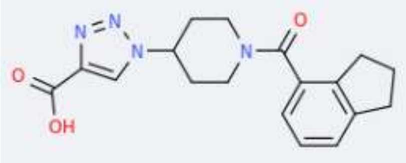 | 340.381 | 0 |

| | | | | | |
|---|---|---|---|---|---|
| 29. | ZINC000262555214 | -12.769 | 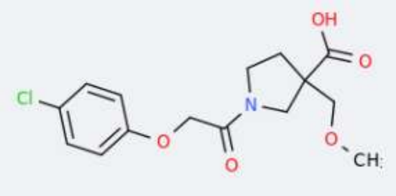 | 327.764 | 0 |
| 30. | ZINC000426580992 | -12.756 | 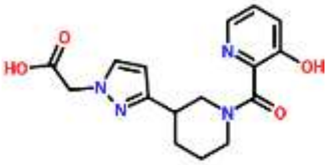 | 330.343 | 0 |
| 31. | ZINC000408587902 | -12.706 | 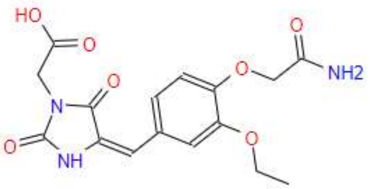 | 363.326 | 0 |
| 32. | ZINC000329072520 | -12.668 | 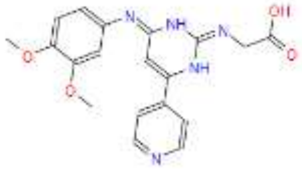 | 381.39 | 0 |
| 33. | ZINC000409124350 | -12.616 | 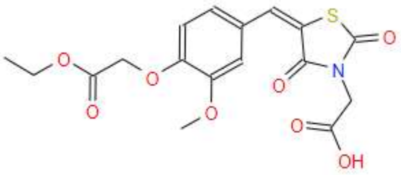 | 395.383 | 0 |
| 34. | ZINC000409133851 | -12.61 | 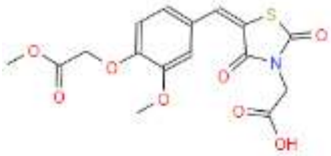 | 381.356 | 0 |
| 35. | ZINC000299751514 | -12.604 | 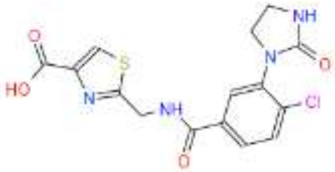 | 380.805 | 0 |
| 36. | ZINC000408592235 | -12.576 | 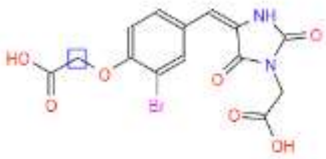 | 399.154 | 0 |

| | | | | | |
|---|---|---|---|---|---|
| 37. | ZINC000666527384 | -12.576 | 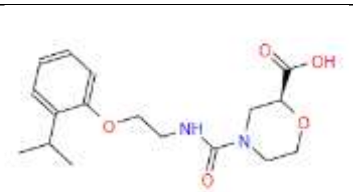 | 336.387 | 0 |
| 38. | ZINC000262311321 | -12.552 | 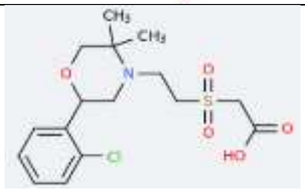 | 375.866 | 0 |
| 39. | ZINC000260680923 | -12.523 | 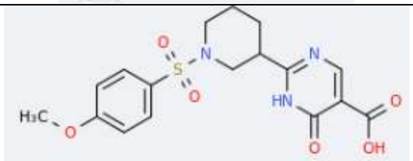 | 393.414 | 0 |
| 40. | ZINC000067139086 | -12.503 | 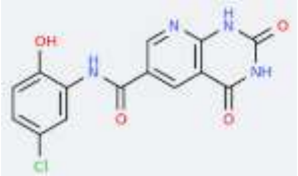 | 332.702 | 0 |
| 41. | ZINC000408589596 | -12.447 | 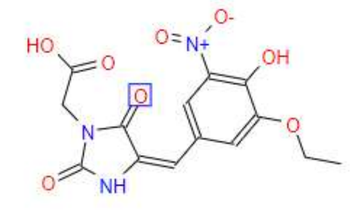 | 351.272 | 1 |
| 42. | ZINC000262416439 | -12.44 | 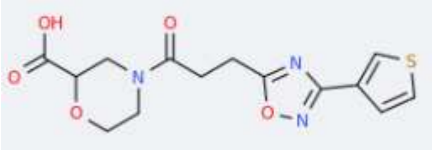 | 337.35 | 0 |
| 43. | ZINC000102754519 | -12.411 | 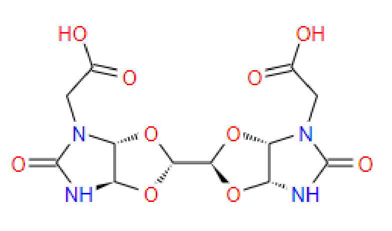 | 374.263 | 2 |
| 44. | ZINC000408589765 | -12.407 | 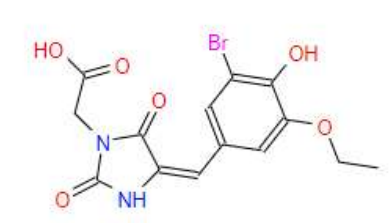 | 385.17 | 0 |

| | | | | | |
|---|---|---|---|---|---|
| 45. | ZINC000328974922 | -12.406 | 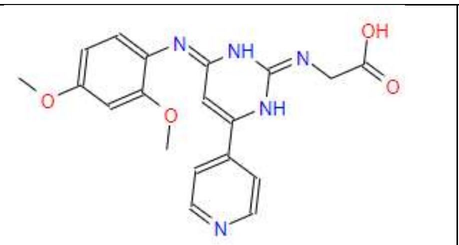 | 381.39 | 0 |
| 46. | ZINC000426503937 | -12.386 | 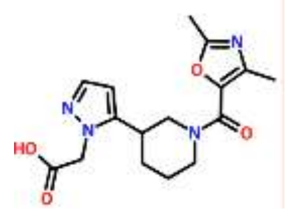 | 332.358 | 0 |
| 47. | ZINC000016027484 | -12.383 | 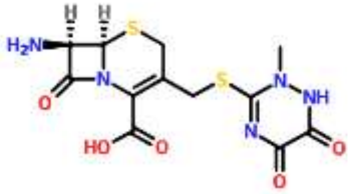 | 371.385 | 0 |
| 48. | ZINC000426505955 | -12.373 | 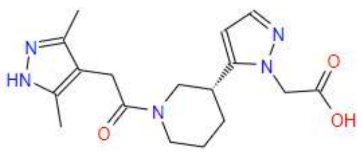 | 345.4 | 0 |
| 49. | ZINC000260677525 | -12.36 | 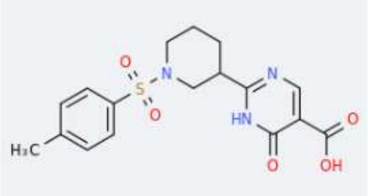 | 377.414 | 0 |
| 50. | ZINC000492118237 | -12.347 | 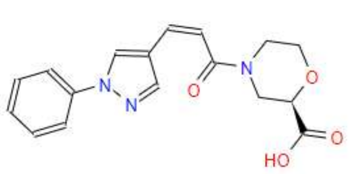 | 327.339 | 0 |
| 51. | ZINC000408593743 | -12.318 | 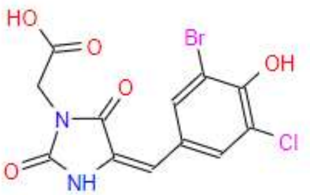 | 375.563 | 0 |
| 52. | ZINC000224969111 | -12.305 | 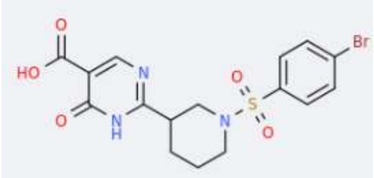 | 442.283 | 0 |

| | | | | | |
|---|---|---|---|---|---|
| 53. | ZINC000408589765 | -12.288 | 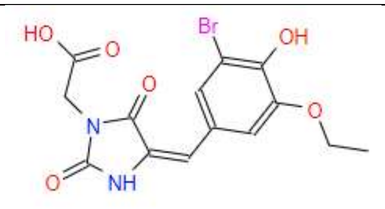 | 385.17 | 0 |
| 54. | ZINC000299747188 | -12.287 | 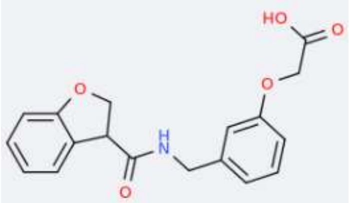 | 327.336 | 0 |
| 55. | ZINC000262846260 | -12.247 | 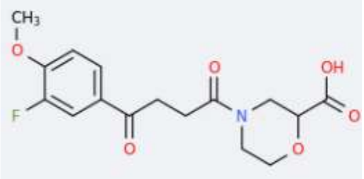 | 339.32 | 0 |
| 56. | ZINC000262974283 | -12.247 | 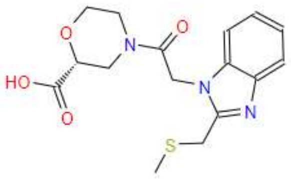 | 349.404 | 0 |
| 57. | ZINC000426668987 | -12.222 | 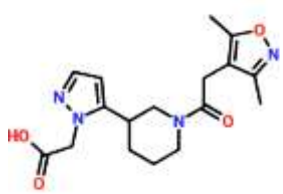 | 346.385 | 0 |
| 58. | ZINC000685944638 | -12.215 | 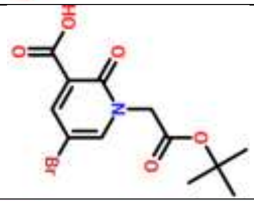 | 332.15 | 0 |
| 59. | ZINC000299748590 | -12.199 | 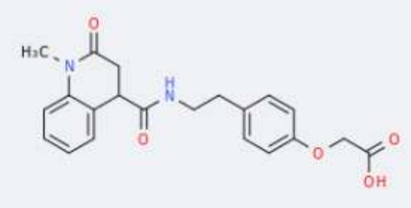 | 382.415 | 0 |
| 60. | ZINC000299751733 | -12.198 | 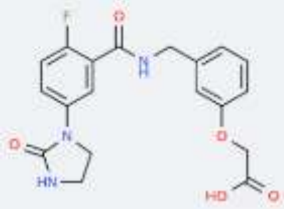 | 387.367 | 0 |

| | | | | | |
|---|---|---|---|---|---|
| 61. | ZINC000329006283 | -12.164 | 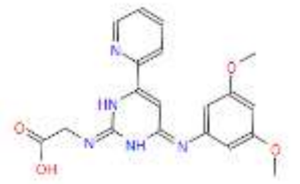 | 381.39 | 0 |
| 62. | ZINC000666527385 | -12.151 | 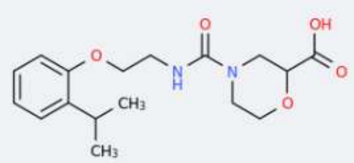 | 336.387 | 0 |
| 63. | ZINC000663050629 | -12.127 | 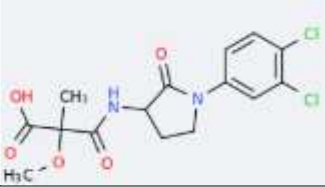 | 375.208 | 0 |
| 64. | ZINC000262843487 | -12.113 | 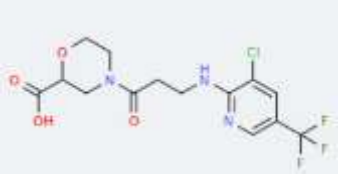 | 381.738 | 0 |
| 65. | ZINC000408595336 | -12.106 | 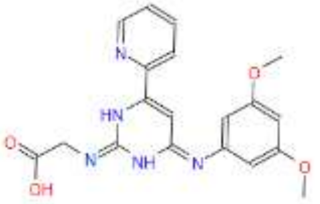 | 392.365 | 0 |
| 66. | ZINC000102621347 | -12.103 | 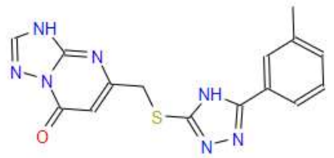 | 339.374 | 0 |
| 67. | ZINC000262949436 | -12.074 | 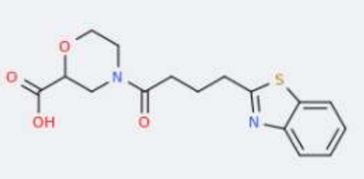 | 334.389 | 0 |
| 68. | ZINC000663218370 | -12.019 | 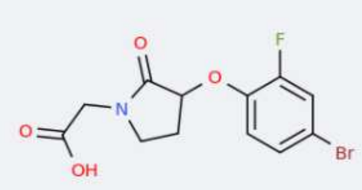 | 332.126 | 0 |

| | | | | | |
|---|---|---|---|---|---|
| 69. | ZINC000005018755 | -11.994 | 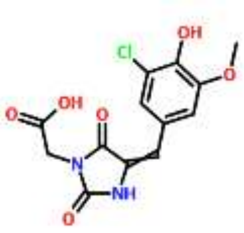 | 326.693 | 0 |
| 70. | ZINC000262906732 | -11.987 | 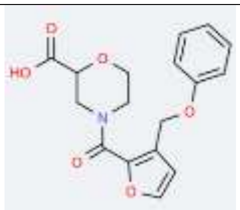 | 331.324 | 0 |
| 71. | ZINC000408596006 | -11.985 | 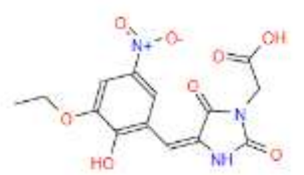 | 351.272 | 1 |
| 72. | ZINC000299749523 | -11.969 | 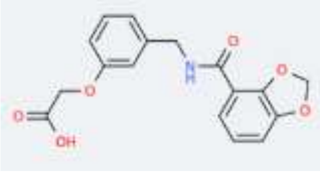 | 329.309 | 0 |
| 73. | ZINC000263398415 | -11.964 | 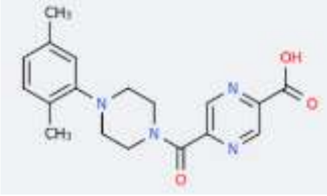 | 340.381 | 0 |
| 74. | ZINC000408594188 | -11.954 | 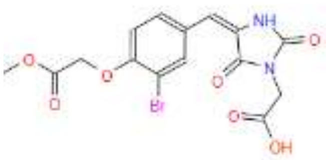 | 413.181 | 0 |
| 75. | ZINC000263008949 | -11.952 | 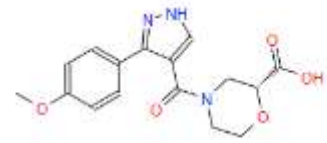 | 331.327 | 0 |
| 76. | ZINC000262895427 | -11.951 | 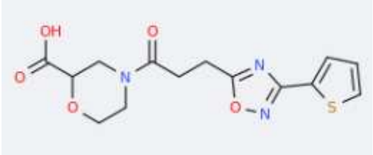 | 337.35 | 0 |

| | | | | | |
|---|---|---|---|---|---|
| 77. | ZINC000408593576 | -11.946 | 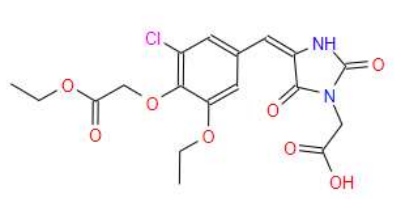 | 426.81 | 0 |
| 78. | ZINC000224969015 | -11.927 | 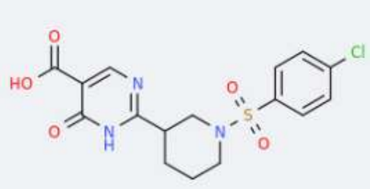 | 397.832 | 0 |
| 79. | ZINC000426491591 | -11.918 | 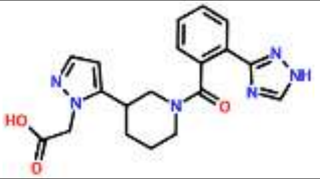 | 380.405 | 0 |
| 80. | ZINC000408589637 | -11.918 | 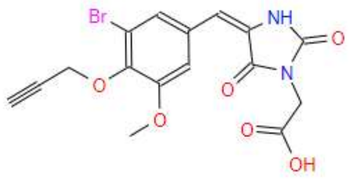 | 409.192 | 0 |
| 81. | ZINC000237456102 | -11.875 | 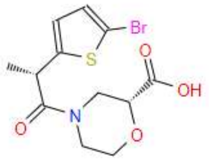 | 348.211 | 0 |
| 82. | ZINC000263005162 | -11.858 | 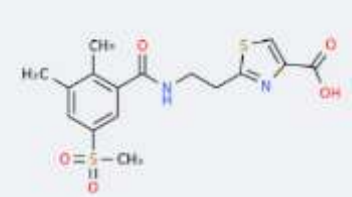 | 382.449 | 0 |
| 83. | ZINC000408590662 | -11.847 | 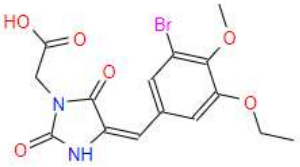 | 399.197 | 0 |
| 84. | ZINC000408591285 | -11.847 | 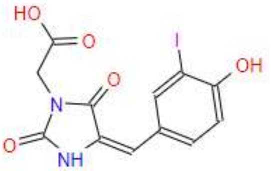 | 388.118 | 0 |

| | | | | | |
|---|---|---|---|---|---|
| 85. | ZINC000299753811 | -11.803 | 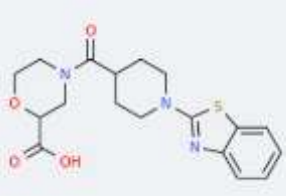 | 375.442 | 0 |
| 86. | ZINC000363215563 | -11.772 | 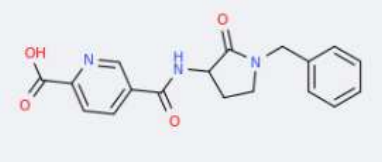 | 339.35 | 0 |
| 87. | ZINC000408513493 | -11.76 | 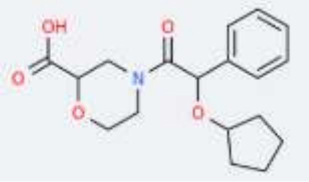 | 333.383 | 0 |
| 88. | ZINC000352226585 | -11.748 | 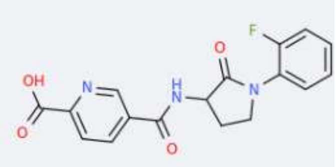 | 343.314 | 0 |
| 89. | ZINC000408593673 | -11.74 | 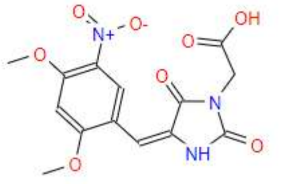 | 375.442 | 0 |
| 90. | ZINC000583650653 | -11.736 | 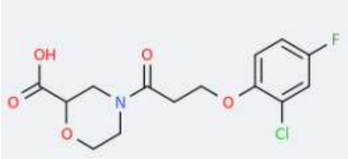 | 339.35 | 0 |
| 91. | ZINC000426515464 | -11.714 | 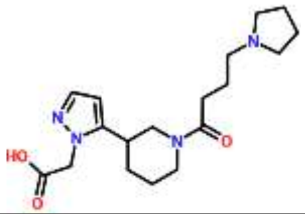 | 333.383 | 0 |
| 92. | ZINC000252383331 | -11.691 | 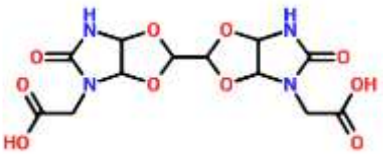 | 343.314 | 0 |

| | | | | | |
|---|---|---|---|---|---|
| 93. | ZINC000237663095 | -11.687 | 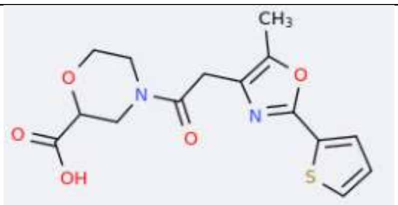 | 351.272 | 1 |
| 94. | ZINC000237965382 | -11.674 | 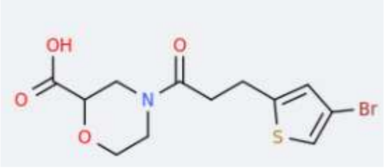 | 331.728 | 0 |
| 95. | ZINC000426512088 | -11.67 | 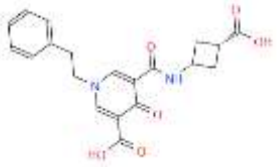 | 348.444 | 0 |
| 96. | ZINC000237965992 | -11.667 | 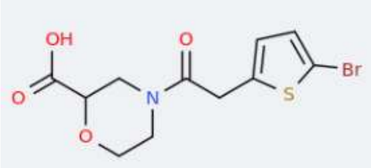 | 374.263 | 2 |
| 97. | ZINC000237966071 | -11.659 | 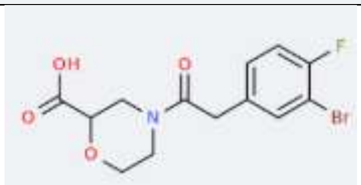 | 336.362 | 0 |
| 98. | ZINC000263008949 | -11.657 | 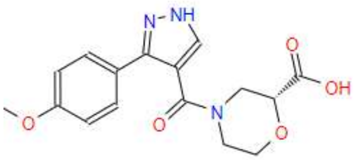 | 348.211 | 0 |
| 99. | ZINC000262715976 | -11.627 | 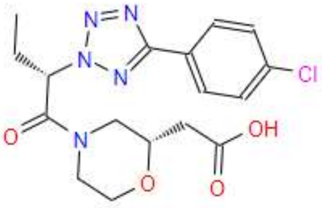 | 384.388 | 0 |
| 100. | ZINC000408592763 | -11.626 | 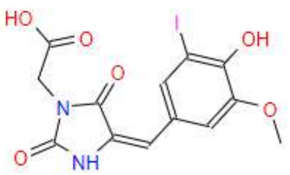 | 334.184 | 0 |

**Note:** In this study, we present the docking scores and Lipinski's rule of five violations for the top 100 small molecules identified as potential Rab10 modulators, as determined using Schrödinger 2024-02 version.

**Supplementary Table S2. ADME/Tox data for the identified top hit compounds for Rab10 with GTP.**

**With GTP:**

| Sample id | Docking score Kcal/mol | Binding energy Kcal/mol | Molecular weight | Donor H-bond | Acceptor H-bond | QPlogPo/w | Percent human oral absorption | Rule of Five violations |
|---|---|---|---|---|---|---|---|---|
| **R1** | -15.46 | -25.41 | 326.36 | 2 | 5.5 | 2.52 | 57.597 | 0 |
| **R2** | -15.24 | -23.14 | 340.13 | 2 | 5.5 | 2.33 | 57.736 | 0 |
| **R3** | -13.99 | -43.21 | 348.78 | 2 | 7 | 1.95 | 63.758 | 0 |
| **R4** | -13.48 | -36.04 | 259.22 | 1 | 5.5 | 3.63 | 83.621 | 0 |
| **R5** | -13.39 | -23.85 | 385.17 | 5 | 9.05 | -1.98 | 19.247 | 1 |
| **GTP** | -20.61 | -43.11 | 523.183 | 5 | 21.60 | -3.38 | 0 | 3 |

**Note:** SwissADME revealed that all the top 5 compounds **R1-R5** have good lipophilicity, are water-soluble, have high gastrointestinal (GI) absorption, and comply with Lipinski's rule.

**Supplementary Table S3. ADME/Tox data for the identified top hit compounds for Rab10 without GTP.**

**Without GTP:**

| Sample id | Docking score Kcal/mol | Binding energy Kcal/mol | Molecular weight | Donor H-bond | Acceptor H-bond | QPlogPo/w | Percent human oral absorption | Rule of Five violations |
|---|---|---|---|---|---|---|---|---|
| **R1** | -11.48 | -14.86 | 326.36 | 2 | 5.5 | 2.52 | 59.05 | 0 |
| **R2** | -11.31 | -26.22 | 340.13 | 2 | 5.5 | 2.35 | 54.93 | 0 |
| **R3** | -7.81 | -45.14 | 348.78 | 1 | 7 | 1.94 | 62.96 | 0 |
| **R4** | -9.73 | -30.79 | 259.22 | 2 | 5.5 | 3.62 | 83.81 | 0 |
| **R5** | -8.68 | -22.30 | 385.17 | 5 | 9.05 | -1.98 | 18.05 | 1 |
| **GTP** | -11.37 | -40.04 | 523.183 | 5 | 21.60 | -3.49 | 0 | 3 |

**Note:** SwissADME revealed that all the top 5 compounds **R1-R5** have good lipophilicity, are water-soluble, have high gastrointestinal (GI) absorption, and comply with Lipinski's rule.

**Supplementary Table S4. Predicted toxicity and pharmacokinetic properties of the top-ranked small molecule modulators (R1–R5).**

| | LD50 (mg/kg) | Toxicity class | Hepatotoxicity | Cardiotoxicity | Carcinogenicity | Immunotoxicity | Mutagenicity | Cytotoxicity | BBB | Neuro-toxicity |
|---|---|---|---|---|---|---|---|---|---|---|
| **R1** | 1000 | 4 | Inactive | Active | Inactive | Inactive | Inactive | Inactive | Inactive | Inactive |

| | | | | | | | | | |
|---|---|---|---|---|---|---|---|---|---|
| **R2** | 500 | 4 | Active | Inactive | Inactive | Active | Inactive | Inactive | Active | Active |
| **R3** | 1190 | 4 | Inactive | Inactive | Inactive | Inactive | Inactive | Inactive | Active | Active |
| **R4** | 1190 | 4 | Inactive | Inactive | Inactive | Inactive | Inactive | Inactive | Active | Active |
| **R5** | 1000 | 4 | Active | Inactive | Inactive | Inactive | Inactive | Inactive | Active | Inactive |

**Note:** Toxicological profiles were evaluated using the ProTox-II prediction platform. All compounds were classified as **Toxicity Class 4**, indicating moderate acute toxicity. The identified compounds possess generally acceptable safety profiles for further experimental validation and optimization.

| Compound | Rab10 (Kcal/mol) | Rab8 (Kcal/mol) | Rab11 (kcal/mol) | Rab5 (Kcal/mol) |
|---|---|---|---|---|
| **R1** | **-10.21** | -3.71 | -5.42 | -13.24 |
| **R2** | **-13.76** | -5.42 | -5.08 | -11.24 |
| **R3** | **-8.1** | -3.64 | -5.18 | -9.05 |
| **R4** | **-12.52** | -2.56 | -5.02 | -11.61 |
| **R5** | **-9.43** | -3.86 | -4.71 | -8.56 |

**Supplementary Table S5.** Predicted binding of **R1–R5** to GDP-bound Rab10 and representative Rab-family proteins. Molecular docking scores (kcal/mol) of compounds **R1–R5** against nucleotide free Rab10 (PDB: 5LPN), Rab5 (PDB: 8XTU), Rab8 (PDB: 4LHY) and Rab11 (PDB: 3BFK). Docking was performed using the same computational workflow with the GDP binding site. More negative docking scores indicate more favorable predicted protein–ligand interactions.
